# Temporal Coherence Shapes Cortical Responses to Speech Mixtures in a Ferret Cocktail Party

**DOI:** 10.1101/2024.05.21.595171

**Authors:** Neha Joshi, Yu Ng, Karran Thakkar, Daniel Duque, Pingbo Yin, Jonathan Fritz, Mounya Elhilali, Shihab Shamma

**Affiliations:** Electrical and Computer Engineering Department, University of Maryland College Park, MD; Electrical and Computer Engineering Department, The Johns Hopkins University, MD; Institute of Neuroscience of Castilla Y León, University of Salamanca; Institute for Systems Research, University of Maryland College Park, MD; New York University, NYC, New York; Départment d’étude cognitives, école normale supérieure, PSL, Paris

**Keywords:** Auditory Stream Segregation, Cocktail Party Problem, Temporal Coherence, Auditory Cortex

## Abstract

Segregation of complex sounds such as speech, music and animal vocalizations as they simultaneously emanate from multiple sources (referred to as the “cocktail party problem”) is a remarkable ability that is common in humans and animals alike. The neural underpinnings of this process have been extensively studied behaviorally and physiologically in non-human animals primarily with simplified sounds (tones and noise sequences). In humans, segregation experiments utilizing more complex speech mixtures are common; but physiological experiments have relied on EEG/MEG/ECoG recordings that sample activity from thousands of neurons, often obscuring the detailed processes that give rise to the observed segregation. The present study combines the insights from animal single-unit physiology with segregation of speech-like mixtures. Ferrets were trained to attend to a female voice and detect a target word, both in presence or absence of a concurrent, equally salient male voice. Single neuron recordings were obtained from primary and secondary ferret auditory cortical fields, as well as frontal cortex. During task performance, representation of the female words became more enhanced relative to those of the (distractor) male in all cortical regions, especially in the higher auditory cortical field. Analysis of the temporal and spectral response characteristics during task performance reveals how speech segregation gradually emerges in the auditory cortex. A computational model evaluated on the same voice mixtures replicates and extends these results to different attentional targets (attention to female or male voices). These findings are consistent with the temporal coherence theory whereby attention to a target voice anchors neural activity in cortical networks hence binding together channels that are coherently temporally-modulated with the target, and ultimately forming a common auditory stream.

## INTRODUCTION

It is still largely a mystery how humans and other animals can effortlessly select and listen to one target source of interest among many in a cluttered scene of simultaneous speakers or other environmental sounds. This problem of Auditory Scene Analysis [1] (also known as the Cocktail Party Problem) has been the subject of extensive studies over decades [2,3]. Physiological experimentation in behaving humans with speech and other complex stimuli using Electroencephalography (EEG), Magnetoencephalography (MEG), or Electrocorticography (ECoG) [4–12] have shed considerable light on the role of selective attention in the segregation process, confirming that neuronal responses to a mixture tend to reflect the focus of attention.

Physiological and psychoacoustic experiments with simpler stimuli have revealed a potentially powerful mechanism at play in accomplishing this segregation process, namely *Temporal Coherence* [13–14]. The premise of this theory is that acoustic features of a single source tend to co-vary together (e.g. a person’s fundamental frequency will co-vary in time with the person’s speech formants or spatial location). As such, neural responses evoked by a single source will be temporally coherently modulated; hence promoting perceptual binding of coherent neural channels by rapidly forming excitatory connections that ultimately link and enhance responses to all the features of a single source. Independent sources are incoherently modulated relative to each other and hence evoke mutually de-synchronized neuronal responses. This is hypothesized to induce suppressive interactions among co-existing streams that compete for eminence. Temporal coherence is postulated to operate in tandem with processes of attention whereby selective attention influences the coherence process by anchoring neural activity (enhanced phase alignment) between distributed neuronal clusters [30]. This causal enhancement boosts the synchronized responses hence resolving competition between different channels and reinforcing the perceptual separation between overlapping sources. Such modulation has been observed in a wide range of sensory modalities including auditory, visual, and somatosensory cortex [31–33].

Recent experiments in behaving animals have provided empirical support of the temporal coherence hypothesis by revealing enhanced versus suppressed neuronal sensitivity following exposure to sequences of synchronized tones (perceived as one stream) versus alternating tones (two streams) [15]. Similarly, EEG and MEG recordings in human subjects switching attention between two competing streams of alternating tone-chords confirmed that the resulting enhancement to the attended stream is pervasive encompassing all synchronized responses of a chord [16,9]. Moreover, a strong validation of the temporal coherence theory with complex multitalker stimuli came from ECoG recordings in human subjects [17]. Neural recordings in nonprimary auditory cortex (superior temporal gyrus, STG) revealed profound segregation effects with neural responses driven primarily by the attended speaker, regardless of the degree of acoustic overlap with the unattended speaker. In contrast, neural sites in primary auditory cortex (Heschl’s gyrus, HG) revealed responses to both talkers with little effects from attentional selection. Interestingly, the selectivity in STG appeared to be mediated by the correlated responses evoked by each speaker independently of the other; providing further evidence that coherent activity across neural sites can be modulated by selective attention to promote segregation between sources in service of behavioral tasks.

One of the limitations of this literature particularly with human subjects is that the broad spatial resolution of EEG, MEG and ECoG recordings does not allow monitoring of single-unit responses as they become simultaneously enhanced or suppressed when attending to a target stream in a mixture of two competing speech streams. Furthermore, it is difficult to truly evaluate causal effects of attentional feedback on activity of individual neurons to gage changes in neural responses that are directly linked to the neuron’s selectivity. The relative ease of recording single-unit responses in animals is countered by the challenge of training them to segregate complex stimuli such as speech. Earlier animal studies of temporal coherence [18,19] did not engage attentional switching between two competing streams, nor did they use streams of more than one constituent spectral component, making it difficult to directly observe coherence-induced changes of neural channels within a stream or possible inhibitory interactions across competing streams.

In the experiments reported here, we trained animals to listen to a two-voice mixture (female and male speakers), attend to the female voice, and react when she utters a specific target word. We recorded single-unit responses in primary (**A1**), dorsal posterior ectosylvian gyrus (**PEG**), and frontal cortex (**FC**). During task performance, representation of the female speech became enhanced relative to that of the (distractor) male in all three areas. Specifically, cortical neurons in A1 whose responses were dominated by the male voice often became relatively suppressed, while those predominantly driven by the female voice became more enhanced. These effects were observed in A1 but were more amplified in PEG. In FC, only the attended female words responses survived. In fact, the segregation began soon after the onset of the speech mixture and was already evident during reference words prior to the final target utterance by the female. At first glance, these results align with the putative role of selective attention, whereby the attended voice (the female) emerges more prominently across the hierarchy of auditory cortical fields, culminating in exclusive activation of the target in frontal cortex. Delving deeper into these effects reveals a causal increase in information flow that modulates female-responsive neurons more effectively than those responding to male voices. This modulation is specifically facilitated by the temporal alignment among channels belonging to the same auditory stream. This observation is supported by a computational model that incorporates the mechanics of temporal coherence and explores effects of different attentional targets to a female versus male voice.

## Results

Two ferrets (**S** & **A**) were trained to segregate two simultaneous (male and female) speakers, and to attend and react only when the female uttered a specific target word. Responses from a total of 337 single units in the auditory cortex of the two animals (**S**: 101 A1, 138 PEG; **A**: 33 A1, 65 PEG) were recorded, while they listened passively (and subsequently, actively) to the individual (male or female) and mixture speech. In addition, we recorded 54 single-units in the FC of ferret (**S**) during passive listening and performance of the segregation task, and 93 single-units in the FC of a third ferret (**C**) during passive and performance of a “female-alone” task in which the ferret detected the female target-word in the female stream but without any distracting male-voice. See **Methods** for stimuli and task structure details.

The animals performed a conditioned avoidance **Go/NoGo** paradigm in which they licked during a variable number (1-6) of female *reference* words and stopped licking immediately upon hearing the female *target* word. The ferrets learned to ignore the simultaneously presented sequence of distracting male-voice words. All words consisted of 3-4 syllables, 0.25 s in duration as shown in **Figure 1A**. Ferret **S** detected the presence of a female target word /Fa-Be-Ku/ at the end of a sequence of the background *reference* words /Fa-Fa-Fa/ (**Fig. 1B**; *1^st^ trace*). Seven different 4-syllabic male words were used as distractors (**Fig. 1B**; *2^nd^ trace*). One in particular - /Fa-Be-Ku- Se/ - is referred to as the *male target-word* because it shared the same 3 syllables as in the female target-word and was frequently presented as a control to ensure that the animal continues to ignore it in favor of detecting the female target. Ferret **A** was trained on different target and reference words. Since the animals were trained to attend to the female-voice, we recorded responses to the passive and behaving *female-alone* conditions, as well as responses to the *mixture* of simultaneous male and female words (**Fig.1B**; 3^rd^ trace). Male words were randomly chosen and played simultaneously with each of the female reference-words, but with different delays (-400ms, -80ms and +200ms) relative to the onsets of the female words (**Fig. 1C**). It was important that the syllables of the two speakers be temporally *incoherent* (persistently *asynchronized*) and hence they roughly alternated (**Fig. 1B**; *bottom trace*). See Methods for details of the stimuli, trial structure, and word sequences.

**Figure 1:**
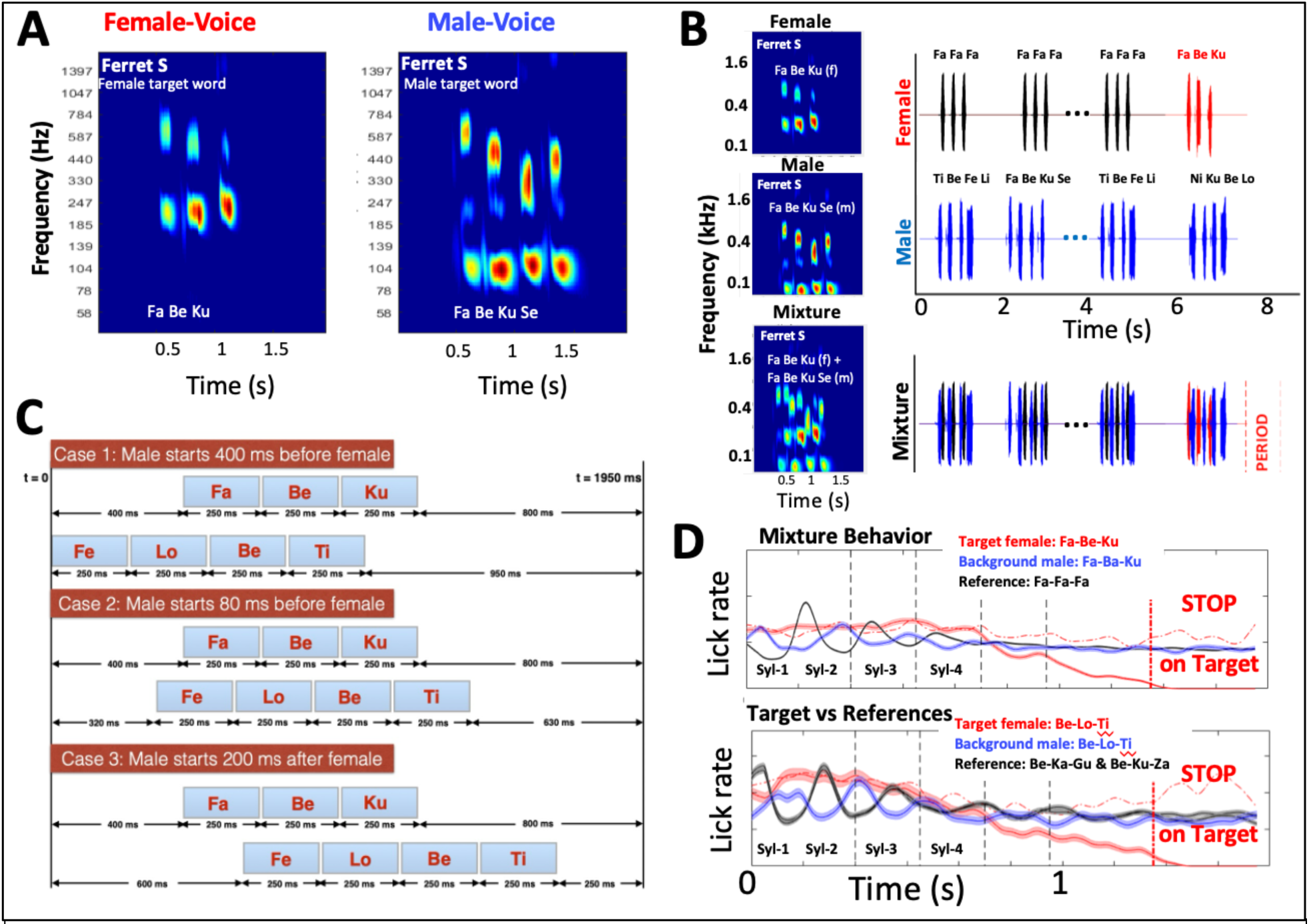
Experiment stimuli, trial structure, and behavioral performance. **(A) Male- and female-voice word spectrograms.** Examples of the target multisyllabic words used in ferret **S** experiments. Female target words consisted of 3 syllables, while the male target words added an extra syllable. **(B) Female/Male streams and mixtures.** The spectrograms display the female target-word (*top panel*), a male word (*middle panel*), and the mixture formed by adding them (*bottom panel*). The structure of the most common trial presentations is schematized by the traces on the right. In the top trace, a female-alone voice stream with reference and target words used in passive presentations, as well as during active detection of the female target-word. The second trace consists of a sequence of various male reference-words that the animals ignore. Finally, the mixture of the words of the two voices. The animal ignores all words in favor of detecting the female target-word. Note that the timings of the male-syllables are always desynchronized (incoherent) from the female-syllables. **(C) Various mixture trials**. The relative timing of the male and female syllables is misaligned to simulate the temporally incoherent sound segments that typically emanate from independent simultaneous talkers. **(D) Lick rates during behavioral performance.** Lick patterns during the various female and male words are depicted in different colors. Lick rates during reference words that the animals ignore remain steady (*black traces*). The animals also ignore the male-words (*blue traces*) and continue licking even when some share all the syllables of the female target-word. Licking stops at the end of the female target-word (*red trace*) as the animals recognize it. The animals stop beyond the red line-marker to avoid a mild shock if they continue.

Finally, **Fig. 1D** demonstrates the behavioral performance of the 2 ferrets (**S** & **A**) that underwent extended neurophysiological recordings during speech segregation. Each of the two panels depicts the lick rates during three critical epochs: (*i*) female-target word in hit *versus* miss trials (red), (*ii*) reference words (black), and (*iii*) male target-word (blue) used as a control. The x-axis depicts time aligned with each of these words in the trials. Ferret **A** (bottom panel) has two black traces corresponding to the two reference words used in the female-voice, while ferret **S** heard one reference female-word. In the case of the single speaker presentation, the black line(s) indicating lick rates to the reference words show a near horizontal trajectory. Contrasted against this is the target-word (hits) licking profile (solid red trace) which indicates that the animal stops licking during the third syllable of the target word presentation. Details of the animals’ behavior are discussed in **Methods**.

### Cortical Representation of Speech Mixtures during Passive Listening

Responses to the speech were recorded in A1 and PEG in ferrets **S** and **A**, as illustrated in **Figure 2A**. Electrode penetrations were made over the two **R**ight and **L**eft auditory cortical hemispheres (***RAC & LAC***) in ferret **S**, and the left hemisphere (***LAC***) in ferret **A**. The tonotopic organization of these fields reveals the approximate presumed extent and borders of the two fields in ferrets **S**(RAC) and **A**(LAC). Recordings were also made in the FC in ferret **S** during task performance and ferrets **S** and **C** during female-alone tasks. Ferrets first listened passively to the speech stimuli of the task and then performed the segregation task afterwards. In the majority of analyzed cells (∼ 95%), evoked responses were phase-locked to the word syllables exhibiting modulations of at- least 2 standard deviations from the baseline neuronal firing rates. **Figure 2B** illustrates the responses to the female target-word in 3 cells that exhibited strong responses to the female-voice (left panel), the male-voice (right panel), or to both (middle panel). We presume that such selectivity in cortical cells in part reflects the coincidence between the cells’ best frequencies (BFs) and the spectral components of the stimuli. To capture this property of the responses, we computed a *speaker selectivity index* (*SSI*) defined as the normalized difference between the variance of the cell’s average male-*alone* response and the variance of the cell’s average female-*alone* target response, both in the passive condition. The SSI ranges between (-1< SSI < 0) for the female-voice preferring cells, and (0 < SSI <1) for the male-voice preferring cells as seen in the 3 examples of **Fig. 2B**.

**Figure 2.**
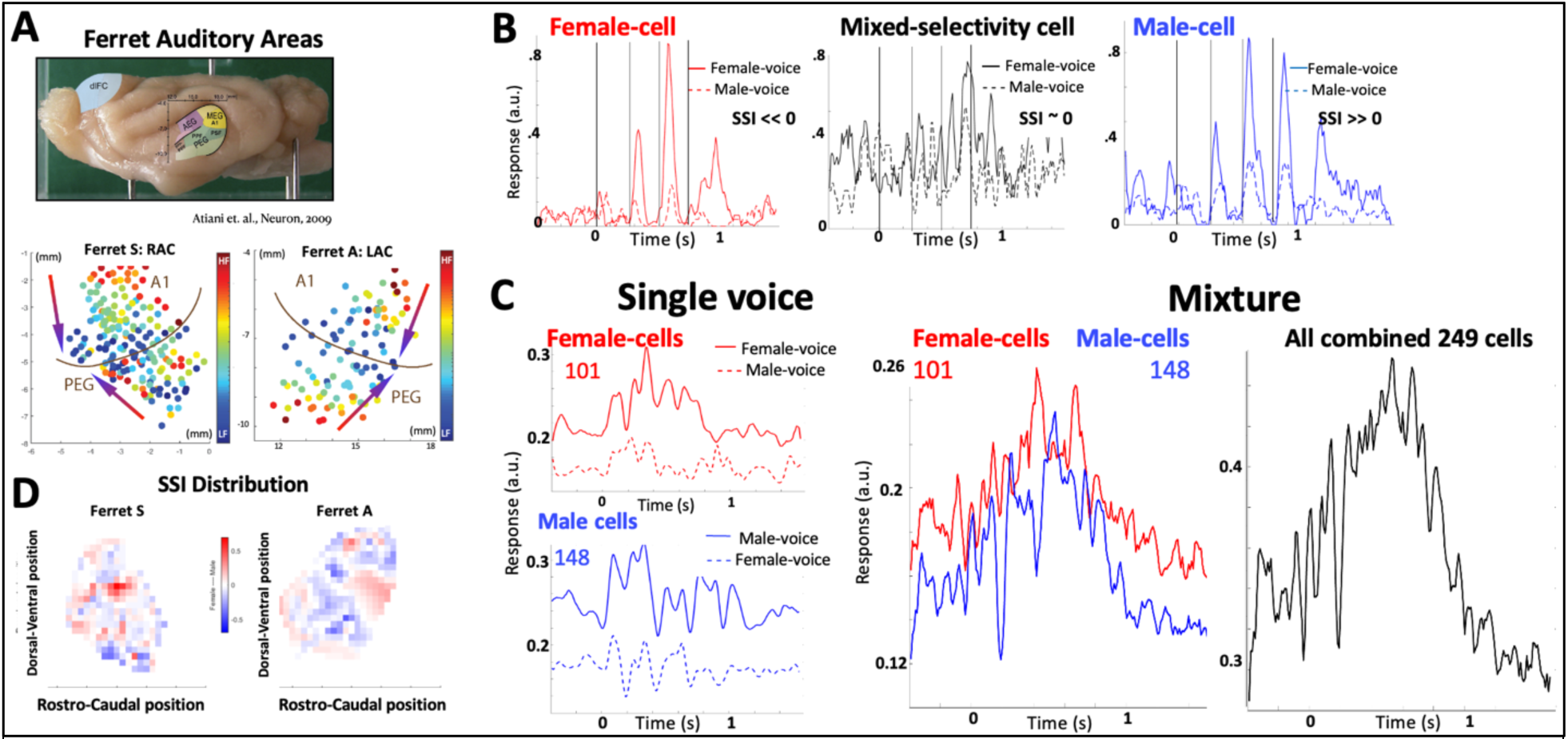
Response properties in the auditory cortex. **(A) Ferret auditory cortex.** It consists of several subdivisions distributed along the anterior, medial, and posterior ectosylvian gyrus (*top panel*). This study focused on responses in the medial and posterior areas of the primary (A1) and secondary auditory cortices (PEG). (*Bottom panels*) The extent and boundaries of these two fields are depicted by the tonotopic organization of high(red)-to-low(blue) BFs in ferrets **S** & **A**. Auditory responses were also recorded from *dorsolateral* Frontal Cortex (dlFC) due to its potential fundamental involvement in complex behaviors such as sound segregation. **(B) Single-unit PSTH responses**. Three auditory cortical cells exhibit diverse responses to the male- and female-voices (blue and red curves, respectively) with strengths that reflect in part the coincidence between their BF’s and the spectral components of the stimuli. The stimulus selectivity index (SSI) characterizes the relative strength of the responses to the two voices, ranging from -1 (sensitive only to female voice) to 1 (sensitive only to male voice). (**C**) **PSTH of population responses to female and male target-words.** Auditory cortical cells in ferret **S** are clustered according to their SSI into female-cells (SSI<0) and male-cells (SSI>0). (*Left panels*) The average responses of these two clusters (101 and 148 cells) to the two voices. When the two voices are presented simultaneously as a mixture (*middle panel*), each cluster responds selectively, in effect segregating the representation of the two voices. However, if the two clusters’ responses are undifferentiated, then the combined PSTH response does not reflect either of the voices or words (*right panel*). **(D) SSI distribution in the auditory cortex** of two ferrets appears scattered and intermingled. It is entirely dependent on the spectral nature of the stimuli, and hence it changes with different voices.

The SSI across all recordings were computed and the cells clustered into two groups that responded completely differently as expected. **Figure 2C** shows responses of a first group, female-cells (*left-top panel*), from ferret **S (**RAC) recordings and reveals a strong average PSTH response to the female-target word (solid) relative to the male-target word (dashed). The second group, referred to as male-cells, exhibits the opposite pattern (*bottom left panel*) with female-target responses weak (dashed) compared to responses to the male-voice (solid). These preferential responses to male/female voices based on neuron’s SSI identity were observed in all individual ferrets and hemispheres **A(**LAC) and **S(**LAC, RAC**) (Suppl. Fig S1)**. Because of this intrinsic response selectivity of various cells, a mixture stimulus containing both the male and female words (right panel) differentially drives the two populations (solid red *versus* blue lines), each reflecting the responses of the spectral components of its selected segregated stimulus.

Note that these segregated PSTH responses are induced in passively listening animals. However, because of the overlapping harmonics and formant transitions of the two voices, combined with the diverse intrinsic selectivity of cortical cells, the female (red)- and male (blue)-cells are rather intermingled and scattered across the ACX (**Fig.2D**). Therefore, speaker segregation of responses based on reading out separately each of these cell groups is (in principle) viable only if the two cell groups remain spatially stable and well-defined. However, it is evident that if the two speakers change their pitches and timbres, then the two cell clusters would change and must be redefined. This is a topic explored further in the computational modeling section.

### Target Enhancement and Distractor Suppression During Task Engagement

The above PSTH of the population responses conceals the profound changes that single-unit responses undergo when the animal engages actively in a segregation task, detecting the female target word amid the distractor male-words. **Figure 3** illustrates effects of rapid plasticity undergone at the population level across female and male cells (clustered based on their SSI). **Figure 3A** depicts the overall clustered responses of all 112 & 98 auditory cortical (ACX) cells in ferrets **S** &**A**. Initially during passive listening, the PSTH levels and modulations of the two clusters are comparable (thin red and blue lines in *top panels*). During task engagement (active), the amplitude of the response *modulations* in the female-cells increases significantly relative to the passive-state, while that of the male-cells diminishes (ferret **S**). This strengthening of the neural response is quantified by the average *variance* of the response in the active and passive states, evaluated for individual neurons in each cluster as shown in **Figure 3B**.

**Figure 3.**
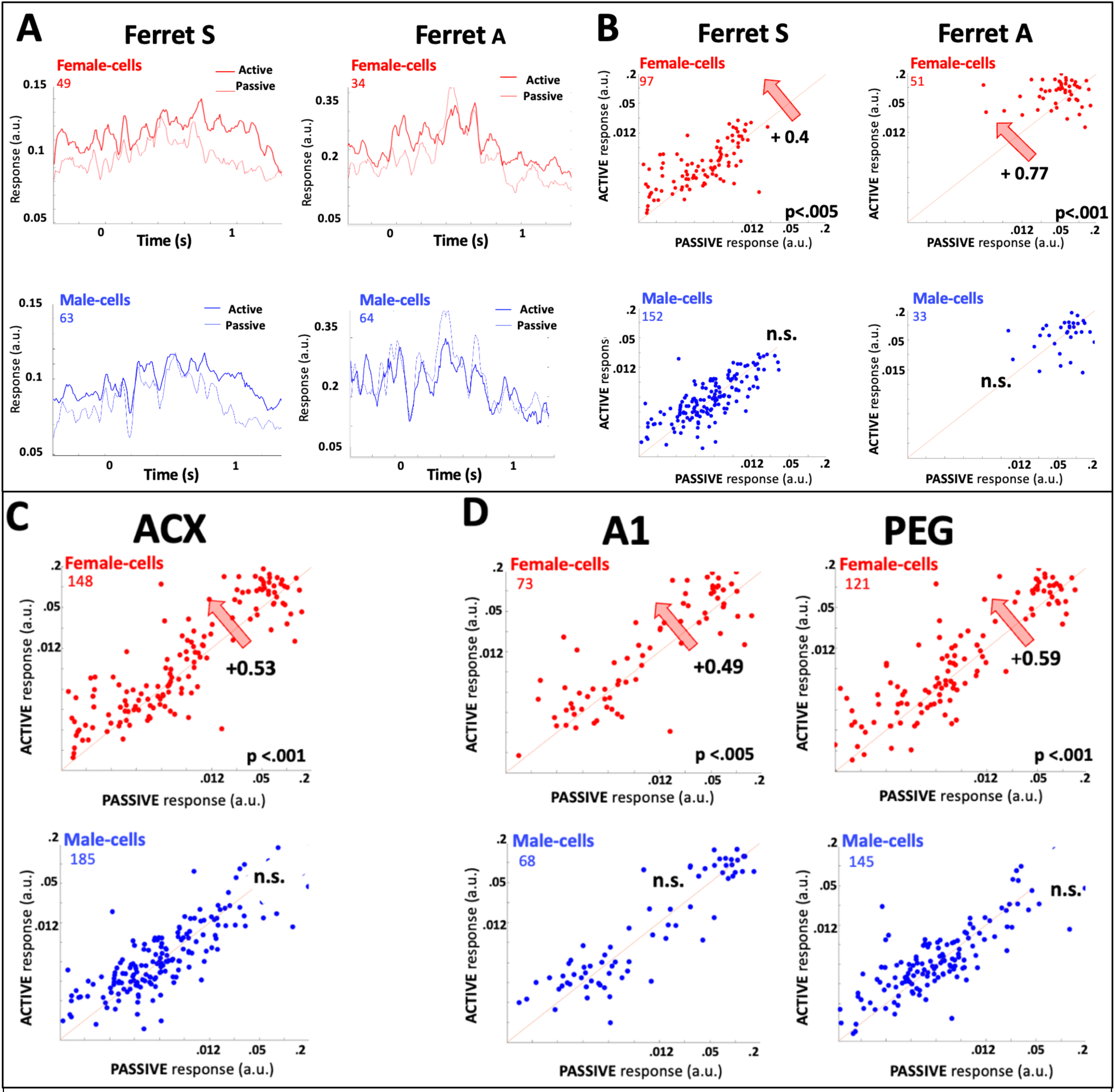
Rapid plasticity of cortical responses during speech segregation. **(A) Plasticity in ACX of ferrets S & A.** During task performance temporally modulated responses of the female-cells are significantly enhanced (*left panels*) relative to those of male-cells which diminish or remain unaffected (*right panel*). **(B) Plasticity in all cells is reflected by the asymmetry of the scatterplots**. Scatter plots depict the changes in the response *variance* of each cell between the passive and active states. The bold numbers in some panels are the overall *effect-size* or extent of asymmetry in the scatterplot (top right numbers) defined as the normalized average of the distances of the points to the midline (see text for details), and the statistical significance of the asymmetry of the distribution of the points around the midline (bottom right number).Note that enhancement in female-cells is the main overall effect in both animals **(C) Plasticity from all cells** recorded reveal a significant female-cell enhancement. (**D) Plasticity in the A1 and PEG**. There is an overall significant enhancement of female cells in both fields. The effects are more profound in PEG than in A1 based on the effect sizes and their significance.

The overall *asymmetry* of the points around the midline captures the *effect-size* of the changes in each population. A signed-distance (+ for above, - for below) of each point relative to the midline is evaluated, *normalized* by the mean of unsigned-distances; resulting in a quantity varying between +1 (enhancement) and -1 (suppression) along with a statistical measure of scatter around the midline (see **Methods**). The scatter of all female-cells in the ACX of both ferrets (**Fig. 3B**) reveals an overall enhancement (*effect-sizes* = +0.4, +0.77) compared to the non-statistically significant global effects in the male-cells. This pattern in the two animals holds if we combine the data from all animals (**Fig. 3C**). Comparing overall changes in fields A1 and PEG of all cells reveals that in A1 (Fig. 3D *left panels*) are relatively smaller and less significant than those in the PEG. Interestingly, examining A1 and PEG patterns for each animal separately reveals a diverse set of effects even in the same animal. For instance, on the one hand, A1 changes are small in ferret **S** but sizable in ferret **A (Fig. S2A).** PEG in ferret **S**(RAC), on the other hand, displays strong suppression of the male-cells, but insignificant effects in female-cells (**Fig. S2B**). The opposite occurs in the PEG of the other two hemispheres (**Fig. S2B**).

### Segregation Viewed Through Stimulus Reconstructions from Auditory Cortex Responses

Looking beyond the PSTH responses and scatter plots, we also examined the responses at the population level to assess their contributions to the details of the speech segregation task in different cortical regions. Here, we employ linear reconstruction of the stimulus, as outlined in [23,24]. Briefly, we reconstruct the spectrogram representations implied by the population activity by learning the *inverse filters* that map the responses to single voice stimuli presented passively (See **Methods**). We then reconstruct the spectrograms implied by the responses while the animal listened to the *mixture passively*, and *actively* performing the segregation task. Comparing the two reconstructed spectrograms reveals the effect of task engagement, and how the auditory cortex extracts and enhances the female-voice relative to the male-distractor. Because of the difference in speech stimuli used with ferrets **S** & **A**, we focus first on the responses in ferret **S (Fig. 4)** and provide analogous results from ferret **A** in **Figure S3.**

The panels in **Figure 4A** are reconstructions of the responses to passive listening of the female/male-alone target words from the entire auditory cortex (A1 & PEG). In all panels, the *blue* and *red* arrows (and dashed lines) mark the frequency channels dominated by the male and female voices, respectively. In the female- and male-alone panels (**Fig. 4A**), the frequency channels corresponding to the first 4 harmonics of each voice are marked by dashed lines. The *red* and *blue* arrows mark the channels mostly driven by the corresponding female- and male-voice. The choice of pitches and formant structure of each voice thus facilitated a clear view of each voice’s responses and the effects of engagement on them.

**Figure 4.**
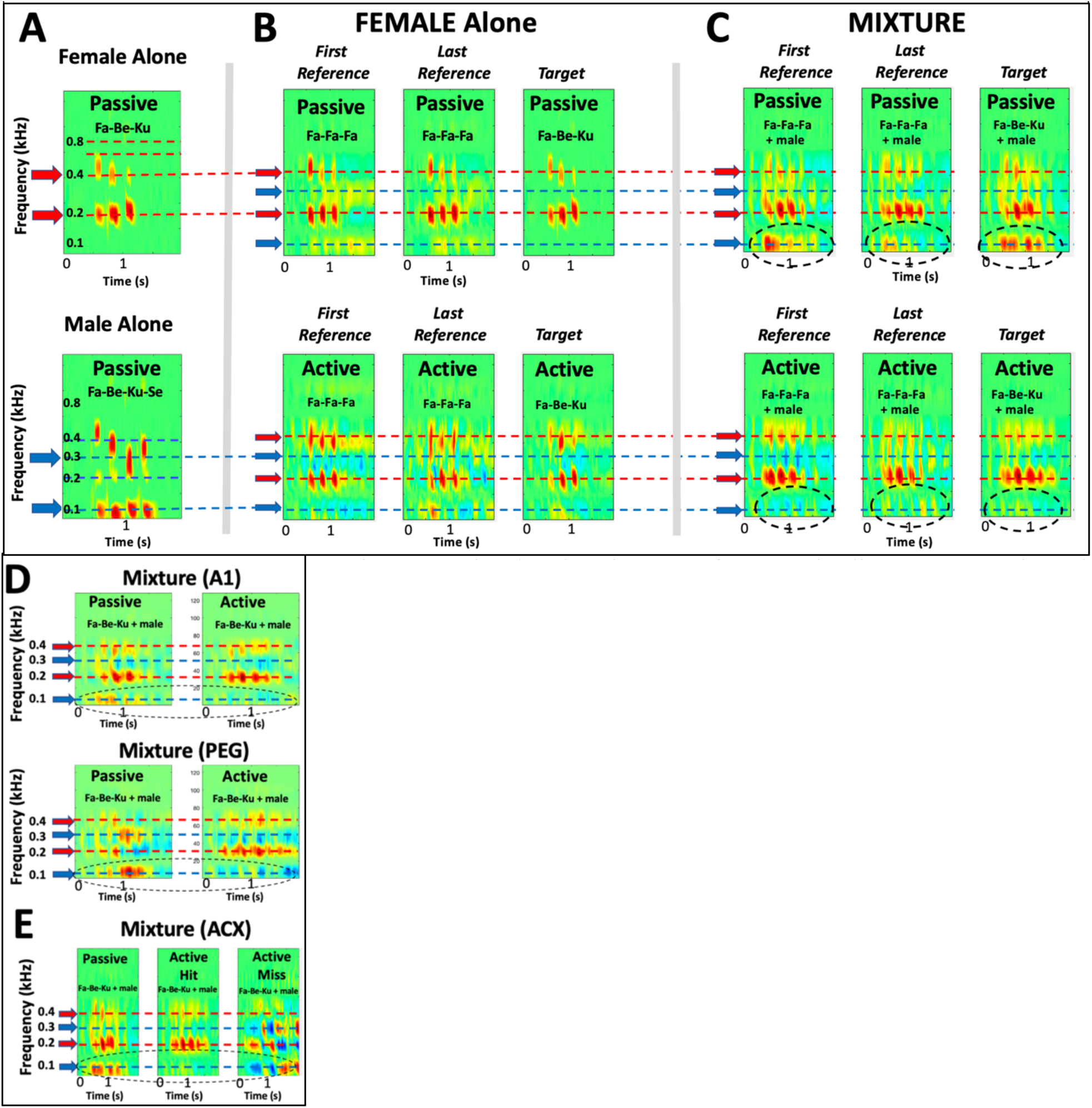
Reconstructions of passive and active spectrograms from ACX of ferret S. **(A) Reconstructions from the single-voice stimuli.** The female and male target-word spectrograms are reconstructed during the passive state. The reconstructions resemble closely the corresponding stimulus spectrograms (Fig. 1). The arrows and dashed lines highlight the frequencies dominated by the male harmonics (*blue*) and the female harmonics (*red*). **(B) Reconstructions from the single female-voice.** The reconstructions are similar during passive (*top panels*) and active (*bottom panels*) states, except for the appearance of weak suppression of the male channels in the active panels. **(C) Reconstructions of the mixture spectrograms.** During the passive state (*top panels*), the reconstructed spectrograms display the spectral features of both voices (activity in the channels of the male- and female-voices, marked by arrows and dashed lines). During the active state (*bottom panels*), the male frequency channels become significantly suppressed in both target and reference words (as highlighted within the dashed ovals). **(D) Reconstructions in A1 and PEG.** Male distractor channels are suppressed in the active (relative to the passive) state in both A1 and PEG (highlighted by the dashed ovals). **(E) Differences between representation of female target words during *hit* and *miss* trials.** In the active state, the suppression of the male-voice channels (blue dashed lines) is complete during the successful hit trials (*middle panel*) but fails significantly during the miss trials (*right panel*).

As expected, the reconstructed spectrograms are very close approximations of the original spectra (*top panels* of **Fig. 1A)**. **Figure 4B** shows the reconstructed spectrograms of the female-alone responses during passive (*top panels*) and active (*bottom panels*) states. The reconstructions shown are of the *first* and *last* of the sequence of reference words /Fa-Fa-Fa/ within a trial and the female target-word /Fa-Be-Ku/. In both passive and active states, the reconstructions obtained are excellent. Note the faint suppression (blue regions) at the male frequency channels only during the active state despite the absence of any male distractor sounds. It is presumed that these are in part because of task engagement which causes an overall suppression of cortical responses, especially in A1 [35]. We shall explore this issue in more detail in a later section when we analyze the responses in the male and female-cells during the reference sequences.

When the male-distractor words are added to the female speaker, the mixture spectrograms become more complex and noisier. For instance, the spectrograms in **Figure 4C** (*top panels*) are reconstructed from the passive responses to the speech mixture of the male added to the female reference and target words. They do not resemble the female-alone spectrograms (*top panels* of **Fig. 4B**), but nevertheless preserve features of the speaker-alone spectra such as the male and female fundamental frequency responses (∼ 0.1 and 0.2 kHz) and remnants of the higher harmonics or formants of both speakers. However, when the animal engages in the segregation task, the reconstructed spectrograms (*bottom panels of* **Fig. 4C**) undergo a significant transformation with the suppression of the male frequency channels (*blue* arrows and dashed lines) relative to those of the female’s (*red* arrows and dashed lines). The suppression is also evident in the reconstructions of the reference words responses in the female stream indicating that the responses to the attended female are already enhanced during the reference period preceding the female target-word. The disappearance of the male distractor responses in the active reconstructions (in contrast to the passive) is highlighted by the dashed ovals in **Fig. 4C**. In the end, the representation of the attended female during the task approaches that of the female-alone panels (**Fig. 4B)**.

**#Fig. 4D** contrasts the differential contribution of the A1 and PEG fields to the segregation process. In humans, it has been shown that attention and task engagement do not significantly modulate responses in the primary auditory cortex [17]. By reconstructing separately from these fields in the passive and active states, it is evident that *both* fields exhibit the relative enhancement and suppression highlighted earlier in **Figs. 3**. Reconstructions in **Fig. 4D** from PEG responses display at a minimum an equivalent persistent suppression of the male frequency channels at ∼0.1 kHz during engagement (blue suppressed responses within the dashed ovals in PEG and A1). **Fig. 4E** compares male suppression during hit trials with the dramatic failure of the distracter suppression when the reconstructions are made from trials in which the animal *missed* the target-word. Finally, spectrogram reconstructions of PEG responses in ferret **A** are shown in **Figure S3**, and they confirm the overall effects in ferret **S**.

### Temporal Coherence in ACX Cells during Speech Segregation

All the above results are based on clustering the cells into male- and female-cells according to their *SSI*, an index that reflects many factors including the tuning and BF of each neuron. Most cells encountered have a mixed selectivity to the two voices and hence a low *SSI* (as evidenced by the crowding of the points near the origin of all scatterplots). It is therefore likely that *SSI-*based clusters are noisy in addition to being quite variable depending on the specifics of the voices encountered. An alternative clustering that remedies some of these difficulties is to label the cells according to whether their ongoing temporally modulated responses are *phase-locked* to those of the female- or male-voices. Therefore, cells can be readily grouped together if they are highly correlated among themselves, and grouped apart if they are not. This is the essence of the temporal coherence principle discussed in the introduction and that will be exploited next.

ACX neurons usually phase-lock to the modulations of their most effective or preferred stimulus, e.g., female-cells in **Figs. 2B, 2C** reflect the phonemic sequences of the female-target words, while male-cells are phase-locked to the male-words. Consequently, the responses of the male- and female-clusters can be readily differentiated by whether they are mutually correlated or not. To perform this clustering, we compute the average covariance between the *passive* responses of all pairs of ACX cells to *mixture* stimuli of female and male target-words. We then compute the singular value decomposition and use the weights of the largest eigenvector to cluster the cells into the male- and female-cell subsets (see **Methods** for details). The *strength* and *sign* of each cell’s weight reflects its relative contribution to the overall phase-locked response and is referred to as the *Correlation Selectivity Index* (***CSI***). It is important to emphasize here that while the SSI is computed from a cell’s response to *single* voices, the CSI is computed based on its response to the voice *mixture*.

**#Figure 5A** compares the average PSTH responses of the CSI- and SSI-based cell clusters to the mixture stimulus. While similar, the two groupings are not identical because the rate response selectivity of a given cell (e.g., due to its BF) may often be incongruent with its ability to phase-lock to the stimulus. The rightmost panel of **Fig. 5A** highlights the rough correspondence between the CSI and SSI indices. Note that for reasons discussed earlier, there are a significant number of cells with low SSI which reduces the correlation between the two groups. **Figure 5B** panels illustrate the PSTHs of the two CSI-based clusters. When distinguished as of A1 or PEG origin in **Figure 5C** panels, the CSI and SSI clusters exhibit similar female-cell enhancement patterns of plasticity as seen earlier in **Fig. 3D**, with strong effects in PEG, and weaker plasticity in A1. We have also seen the same variability in the plasticity effects in ferret **S**(RAC) which in fact reveals a strong suppression of the male-cells during the active state, similar to that of **Fig. S2B**.

**Figure 5.**
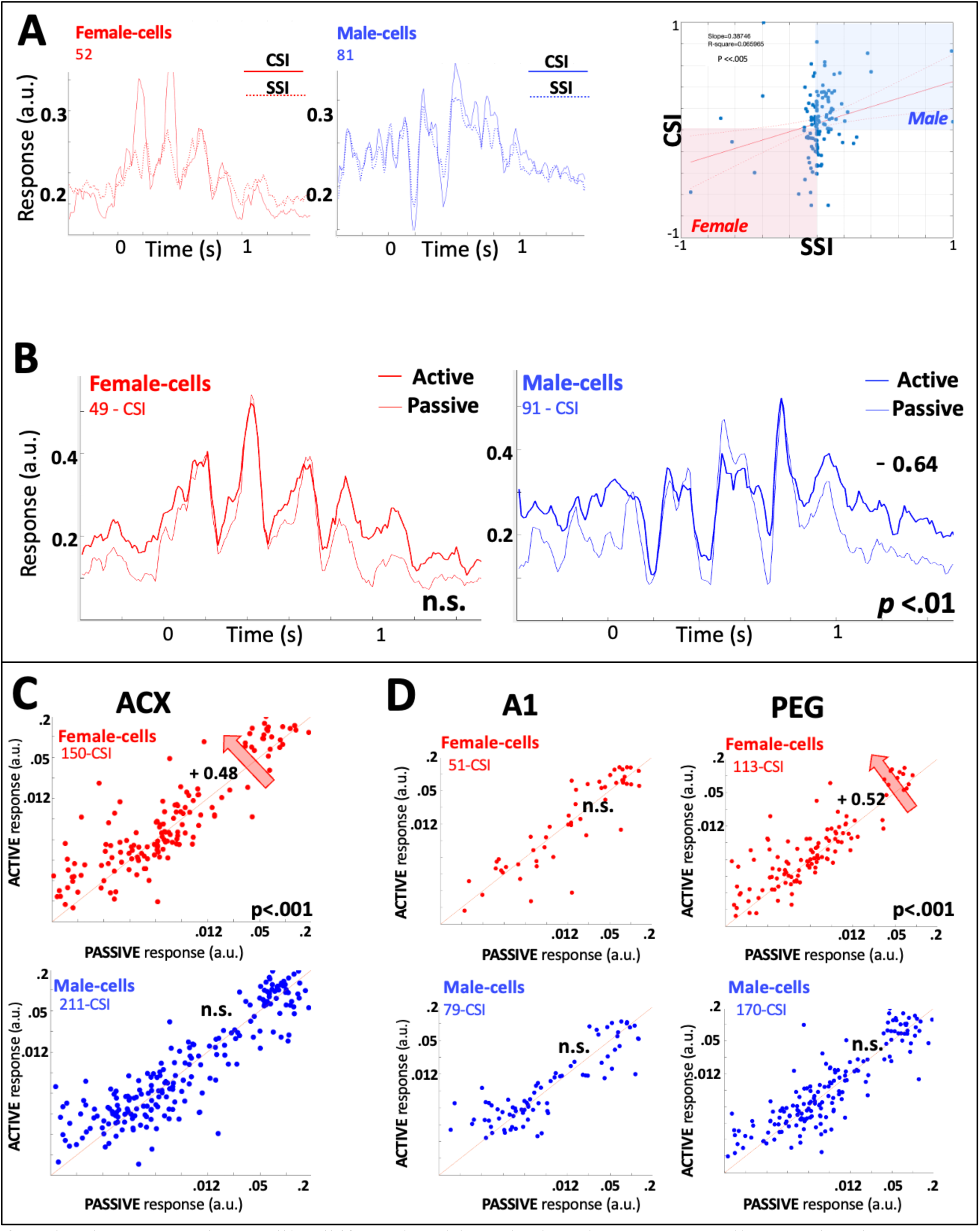
Plasticity in CSI-clustered cells in ferret S(RAC). **(A) Comparing PSTH responses of SSI- and CSI-based clusters**. (*Left and middle panels*) PSTH responses of the CSI and SSI cell clusters to the female-target word (*left panel*) and male-target word (*right panel*) resemble each other, though are not identical reflecting the difference between the temporal and spectral clustering criteria. The scatterplot (right panel) compares the SSI and CSI of all male- and female-cells selected based on their SSI. There is a moderate correspondence between the two populations. **(B) Passive versus active PSTH** responses of the CSI-clusters exhibit similar plasticity patterns as in the earlier SSI-defined clusters of Fig. 3. **(C) Plasticity during active engagement** in speech segregation in A1 and PEG demonstrates strong suppression of the male-cells, thus enhancing the representation of the target female-voice relative to the male distractor. The panels depict the response changes through PSTH and scatterplots exactly as in previous figures. **(D) Scatterplot of all the female and male-cells** aggregated from both animals comparing their responses in passive and active states. Overall effects are like earlier SSI-based results (Fig. 3), with enhancement of the female-cells relative to the male-cells.

### Response Plasticity to the Female-Reference Words

During performance of all tasks, animals must detect in each trial the female target-word (e.g., /Fa- Be-Ku/ in ferret **S**) appearing at the end of a sequence of reference-words (e.g., /Fa-Fa-Fa/) in the female voice. With mixtures, the added male distractor-words must be ignored throughout. We have so far focused on the representation of the responses to the female-target word. The reference words however play an important perceptual role as they form the “female stream” or sequence that the animal must attend to and segregate. If the animal fails to do so, it is likely to miss the female target-word. Here we explore the representation of these reference-words, beginning with the *female-alone* responses in **Figure 6** because of their simpler-to-interpret PSTHs. We then compare the response patterns to the case of the mixture stimuli (**Fig. 6B**). We also provide a schematic (**Fig. 6C**) that summarizes and simplifies the presentation of the complex response changes seen during the sequences.

**Figure 6.**
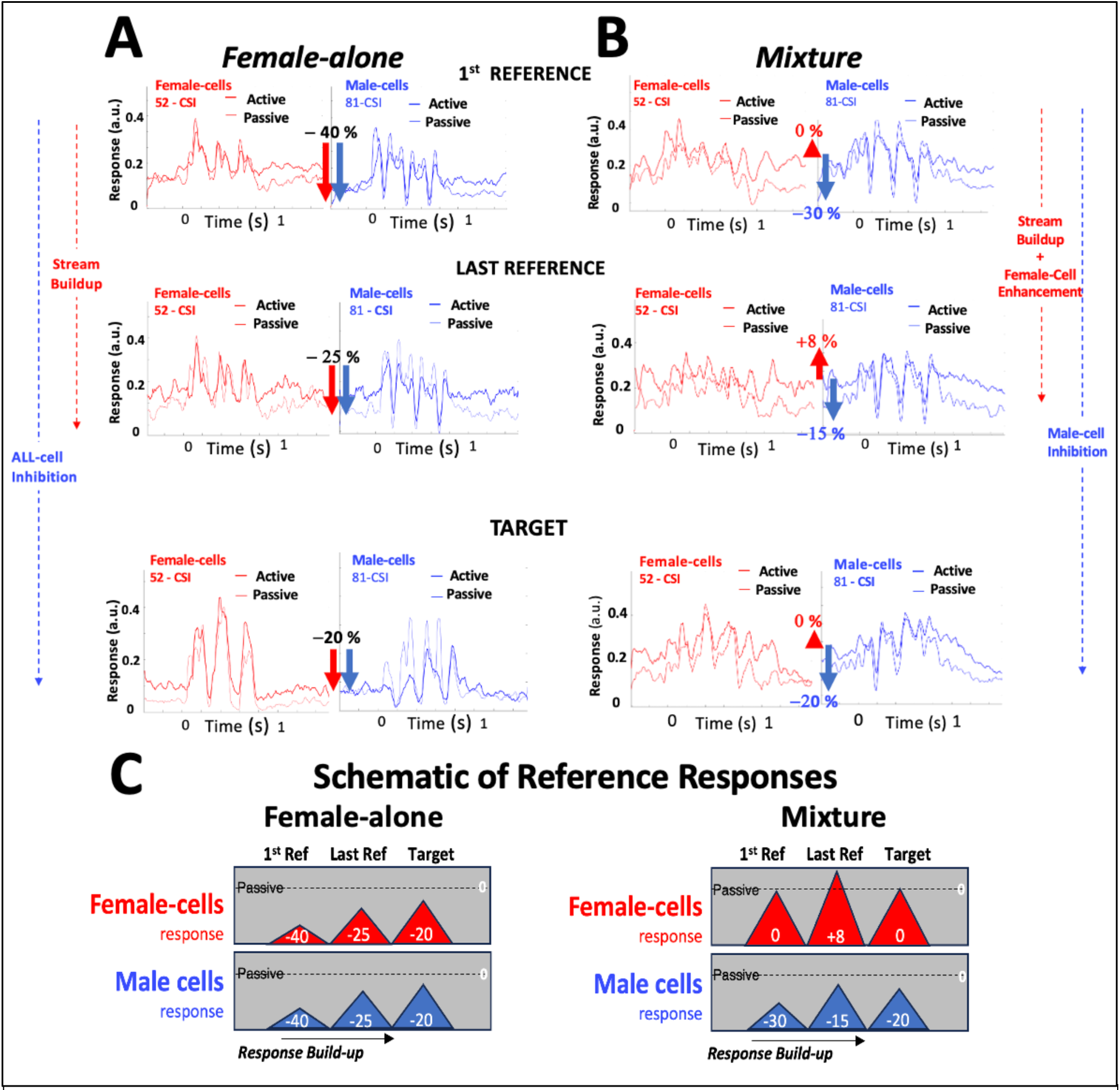
Reference and target responses in female-alone and mixture tasks. **(A) Responses during the female-alone task.** Both male- and female-cells become strongly suppressed (-40%) during the task relative to the passive. Curiously, while the female-cells are phase-locked to the female voice, the male-cells exhibit an *inverted* phase-locking. Responses in both clusters gradually buildup reflecting the formation of the attended female-stream towards the last-reference and target stimuli**. (B) Response changes in the mixture segregation** task are quite different reflecting the *competition* between the two clusters. Female-cell responses remain enhanced throughout compared to the male-cells. Responses again show the buildup of the streams from 1^st^ to last reference. **(C) Schematics of the two task responses.** Triangles represent the responses to the references in female- (red) and male-cells (blue) throughout the trials, relative to the passive levels. During the female-alone task (*left panel*), all responses evolve similarly. During the segregation task (right panel), female-cells remain enhanced throughout compared to the suppressed male-cells. the enhancement of the attended female-voice responses and the suppression of the male-cells.

The evolution of the responses to auditory sequences has been the focus of numerous previous studies in the context of stream formation and temporal coherences [18], in the formation of implicit memory [25], in adaptive efficient coding [26], and in stream formation [28]. A common finding is that sequence responses (especially in the attentive animal and in higher auditory fields) exhibit a *gradual* change following their onset referred to as the “build-up” period of the stream. In our experiments, responses to the reference sequence exhibit a build-up from the 1^st^ to the last reference, as well as the enhancements and suppression seen earlier for the target responses.

For instance, **Figure 6** depicts responses to the *female-alone* sequences during the passive and active states. In the passive state, reference responses of either male- or female-clusters do not experience any significant changes throughout the sequence from 1^st^ to last reference as shown in **Fig. S4**. But upon task engagement, all reference responses become significantly depressed to the 1^st^ reference (−40% relative to the passive response level, as schematized by the red and blue arrows in **Fig. 6A**). All cell responses gradually *buildup* towards the last-reference (-25%), and finally the female target-word (-20%). Curiously, while the female and male responses are roughly equal in power, they are of *opposite polarity*.

During the mixture segregation task (**Fig. 6B**), the responses are dramatically different. *First*, during the 1^st^ reference stimulus, only the male-cells are suppressed (-30%) while the female-cells maintain their response power. The female-voice therefore becomes significantly better represented relative to the male. *Second*, both cluster responses exhibit a build-up towards the last-reference, but with the female-cell cluster exceeding passive levels. *Finally*, we reach the target stimulus where responses maintain the superior level for the attended female-cells relative to the distractor male-cells.

### Speech Segregation in the Frontal Cortex

Ferret FC responds vigorously to auditory stimuli when the animal is engaged in tasks requiring auditory attention as in the speech segregation task. However, FC responses do not normally phase-lock well to their stimuli, making it difficult for instance to reconstruct stimulus spectrograms to identify the source of the responses; instead, we must rely on distinctive features of their PSTHs.

FC responses were recorded in ferrets **S** (54 cells) & **C** (93 cells) during passive listening to the same mixture and female-alone stimuli (**Fig. 1**). Recordings were also made while both ferrets performed the female-alone task. Ferret **S** in addition performed the mixture segregation task. During passive listening, the PSTHs sometimes reflected the stimuli, exhibiting the syllabic structure of the female or male voices when played alone. **Figure 7A** illustrates the average PSTH during passive listening in ferret **S** to the female target-word /Fa-Be-Ku/ (*middle panel*), with the 3 syllabic peaks moderately expressed and aligned to the peaks of the spectrograms (*top panel*). When the animal actively detects the female target-word, the expression of the syllabic peaks of the female target improves considerably (**Fig. 7A**; *bottom panel*).

**Figure 7.**
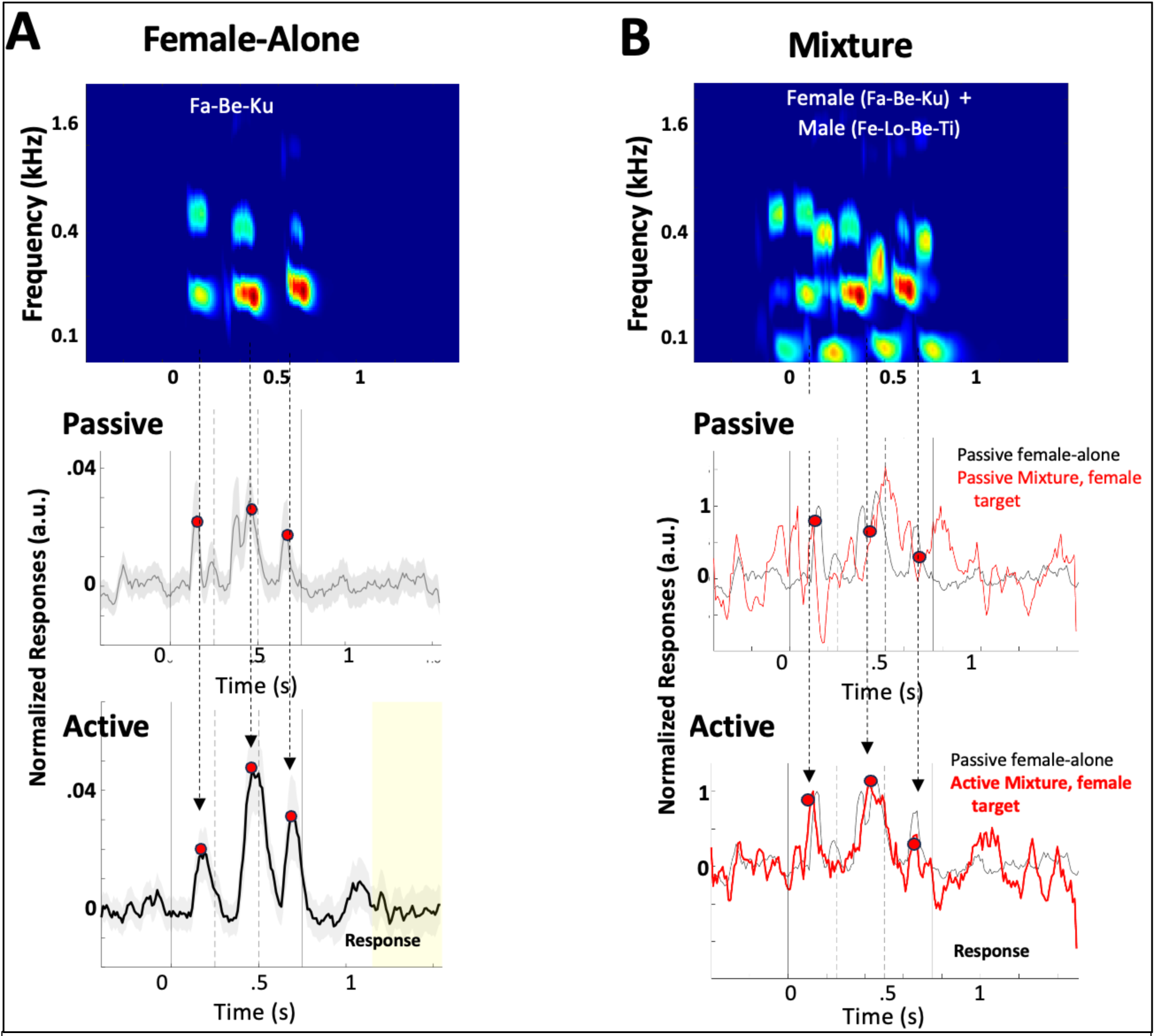
Responses in the frontal cortex in passive and active states. **(A) Plasticity of Female-alone target-word responses.** Stimulus spectrogram (*top panel)*, and corresponding responses in the passive state (*middle panel*), and in the active state (*bottom panel*). The syllabic response peaks (marked by the red circles) of the female target-word become enhanced when the animal engages in the active task. **(B) Plasticity of Female target-word during mixture segregation.** Stimulus spectrogram of target + male distractor words (*top panel)*, and corresponding responses in the passive state (*middle panel*), and in the active state (*bottom panel*). The syllabic response peaks (marked by the red circles) of the female target-word (*thin* black curve) are nearly eviscerated by the additional male-distractor peaks. The target-word responses (*bold* red curve) are enhanced in the PSTH when the animal engages in the active task and begin to resemble the original female target-word (*thin* black curve).

During passive listening to the mixture stimuli, the female target-word induces PSTHs with poorly expressed peaks that do not align with the clean female-alone target-word, as evidenced by the mismatch between the thin red and black curves in **Figure 7B** (*middle panel*). However, when the animal attends to the female target-word during active segregation, the male (distractor) responses are suppressed and the active mixture PSTH changes to resemble closely the shape and peak-alignment of the female-alone target-word, as demonstrated by comparing the thin-black and bold-red curves in *bottom* panel of **Fig. 7B**.

Responses in both ferrets **S** and **C** to the female reference-words in the passive and active states in both the female-alone and mixture segregation tasks recapitulate the findings from the target-word case as seen in **Figure S5**. Finally, note that FC responses in all these analyses are *not* clustered into male- and female-cell responses as they were in the auditory cortex and hence the divergent effects in these two clusters are integrated here.

### Computational account of the speech segregation

To provide an overall framework for the speech segregation process and to further probe the role of temporal coherence in it, we implemented a computational model inspired by the Explicit Memory Multi-resolution Adaptive (EMMA) framework [34]. Like the ferret experiments, the model takes as input a voice mixture alongside a ‘directive’ indicating which stream to be extracted, i.e., an attentional focus similar to how the ferrets were trained to direct attention to the female voice. The model aims to segregate a voice that best aligns with its attentional focus, leveraging two key principles (**Fig. 8A**): **(i)** It maps the input mixture to a nonlinear high-dimensional space mimicking the diverse selectivity of cortical neurons to frequency, pitch, location, and other sound attributes [37] (analogous to the random distributions of **Fig. 2D**). This feature analysis stage is implemented with deep-learning neural embeddings [34], referred to as the *pre-attentional model embeddings* (*Mp*). It is a complex representation that preserves all characteristics of the input mixture as depicted by the mix of female (red) and male (blue) activations in **Fig. 8A**. **(ii)** The responses of the *Mp* stage are next transformed according to the selective attentional focus on the female or male voice. The attentional stage (*Ma* in **Fig. 8A**) achieves this by gating the *Mp* embeddings according to their temporal coherence with the attended target voice, while relegating others to the background. It is implemented in **Fig. 8A** by a projection relative to a phase-alignment with the target [34]. All model parameters illustrated next selected using the same speech material used to train the ferrets (see **Methods**).

**Figure 8.**
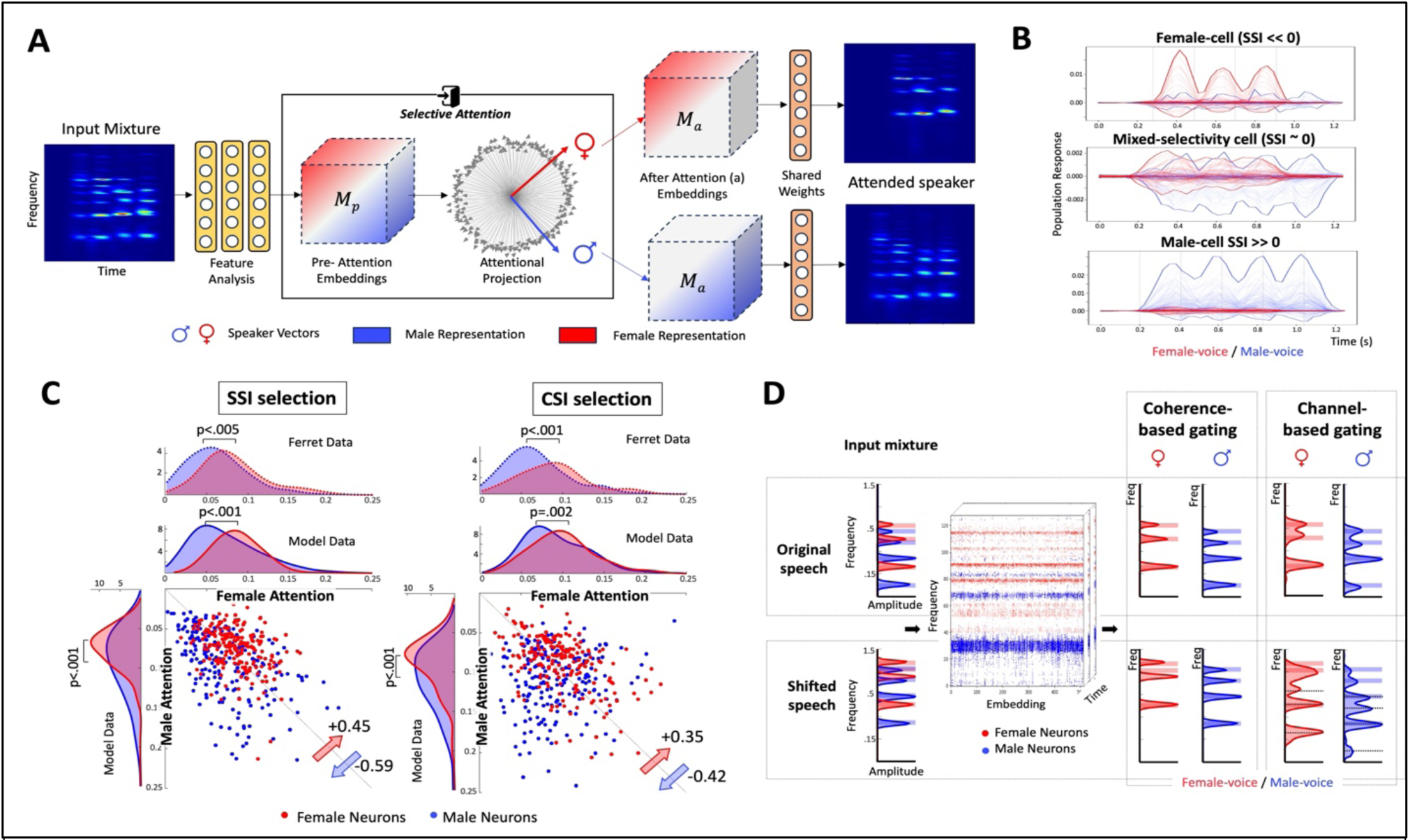
**A. Computational model for temporal coherence in speech segregation**: An input mixture is first analyzed through a multilayer CNN resulting in high dimensional embeddings labeled *Mp* (Model pre-attention). Before attentional selection, the embeddings contain all details of both male and female voices (depicted as a mix of red and blue colors) and span a range of diversely misaligned temporal modulations (or response waveforms with many phases) reflecting the complexity of the input mixture. These are depicted as vectors with different orientations. Depending on the target voice to be attended to, a phase alignment (represented as vector projections) enhances model neuron responses that are temporally coherent with the attended voice (e.g., female) while suppressing others. The selected embedding (referred to as *Ma*) are then mapped back onto a spectrogram of the attended speaker (male or female voice) through a simple inverse transformation. **B. Activations in the feature analysis stage** (*Mp*). In response to single speaker stimuli, some neurons respond best to female- (*top*) or male- (*bottom*), or exhibit a mixed-selectivity (*center*). Therefore, model neurons can thus be clustered according to SSI- or CSI-based indices (e.g., Figs. 2 & 5). Dashed vertical lines represent onsets of the female (*left*) and male (right) syllables, highlighting the phase-locked responses of the model neurons. **C. Mutual information (MI) of pre- and post-attention model responses.** Each scatterplot contrasts the MI between pre-attention and post-attention responses of female-neurons (red points) and male-neurons (blue points) when attending to a female (x-axis) *versus* a male (y-axis) voice. Left (Right) scatterplots represent neuron selection using SSI (CSI) indices, respectively (details in **Methods**). Marginal distributions illustrate the contrast between pre/post-attention MI in male *versus* female neurons when attending to the male voice (y-axis marginal) compared to the female voice (x-axis marginal). Analysis of recorded ferret cortical responses (available only for selective attention to female) is shown above the model marginal contrasting responses of female- and male-neurons. **D. Model responses to original and pitch-shifted speech**. A male/female mixture (averaged spectral profiles shown) is analyzed through the segregation model. Model selectivity to male (blue points) and female (red points) voices (based on SSI) is strongly aligned with their frequency content. This channel-based alignment however is destroyed if the input spectra are shifted (*bottom row of panels*). By contrast, coherence-based gating reliably preserves the representation of the spectral harmonics of the attended male or female voice (as indicated by the dashed black lines in the *top row panels*), even when the pitch of the input voices is shifted (*bottom row of panels*) relative to their original positions.

Attention in the ferrets and the model segregates mixture responses and enhances a target representation relative to the distractor. Before attentional gating (equivalent to passive listening in the ferret) model neurons exhibit a range of phase-locked responses (depicted as continuous activations analogous to neural PSTHs), with some favoring the male voice and others the female or neither. As in the ferret (**Fig. 2B**), model responses can be clustered into male and female-neurons based on their SSI with single-voice responses (**Fig. 8B**), or on CSI with mixture responses (**Figs. 5**). As in the ferret during active engagement, attention in the model *gates* this diversity of responses to segregate and select the appropriate subsets for enhancement and suppression. To quantify this effect, we use the change in *mutual information* (MI) between activations before (*Mp*) and after attention (*Ma*), separately computed in male *versus* female neurons. The primary reason for using the MI measure is that model activations are not strictly positive, and hence response “amplitude changes” alone are not sufficiently informative. MI instead effectively quantifies the degree of certainty before and after attention, analogously to the evaluation of enhancement and suppression based on response variance of the firing rates in the ferret.

**Figure 8C** illustrates the range of MI for neurons selected based on SSI (*left*) or CSI (*right*), comparing effects of attending to the female voice (x-axis) versus attending to the male voice (y-axis) in the same neuron. The analysis distinguishes male (blue points) from female neurons (red points) in both cases. A measure of the effect-size (as in **Figs. 3** & **5**) confirms that when attending to the female voice, coherence-based attentional gating favors responses of female neurons. The opposite effects are seen when attending to a male voice (female neurons - SSI effect-size = +0.45, p<0.001; CSI effect-size = +0.35, p<0.001 - red arrow in **Fig. 8C**). The shifts around the midline reverse in male neurons where attending to a male voice exhibits a stronger predictive pattern (male neurons - SSI effect-size = -0.59, p<0.001; CSI effect-size = -0.42, p<0.001 - blue arrow in **Fig. 8C**). The marginal distributions contrast the effects of attention to the female *versus* male voice in the two groups of (male/female) neurons. They confirm that attention to the female drives female neurons to exhibit a statistically significantly stronger predictive effects relative to male neurons (female attention - SSI, Mann-Whitney U-test, p<0.001; CSI Mann-Whitney U-test, p=0.02 - marginal along the x-axis in **Fig. 8c**). As expected, the opposite effects are observed when attending to the male voice, as male neurons display statistically significant stronger effects than female neurons (male attention - SSI, Mann-Whitney U-test, p<0.001; CSI Mann-Whitney U-test, p<0.001 - marginal along the y-axis in **Fig. 8c**).

We also analyzed all responses recorded from ferrets **S** & **A** using the same MI measure and found that the same trends are observed when the ferrets attended to the female voice. Thus, female-cells consistently showed higher MI values relative to male neurons (SSI, Mann-Whitney U-test, p<0.001; CSI Mann-Whitney U-test, p<0.005 – dashed lines / panel above the x-axis marginal in **Fig. 8c**). All trends were observed consistently *regardless* of the SSI or CSI clustering method. Model results reported above included a selection of 200 female and 200 male neurons (based on both SSI or CSI). We have confirmed that the trends reported are maintained with varying numbers of model neurons, though statistical power starts decreasing when less than 50 neurons are used in the analysis.

Finally, a key conceptual question addressed in this study is the ultimate benefit of temporal coherence as an attentional gating mechanism. An alternative hypothesis is a channel-selection view where the brain relies on feature selectivity prominent throughout the auditory system (e.g. *best frequencies*) to gate channels that are commensurate with the expected target. These two approaches were the motivation for exploring CSI versus SSI based clustering earlier, and they are simulated in **Fig. 8D** where we contrast explicitly the two scenarios: *coherence-based gating* where the pre-attention responses (*Mp*) are transformed using phase-projections (or temporal coherence); and *feature-based selection* where channels responsive to specific features of the target voice are selected and enhanced relative to the others (e.g., *only* female neurons are selected when attending to the female voice and vice versa). The *top row panels* of **Figure 8D** show that this is a plausible alternative if (as in this case) neurons are labeled *apriori* as to their feature selectivity. Thus, when the model attends to one voice in a mixture (average spectral profiles shown on the left to highlight peaks from either voice), it can extract the rightmost spectra either according to the coherence-based gating (*left*) or channel-selection (*right*). While both scenarios are validated here, channel-selection would fail if the spectra were altered such that a different set of features are activated by each voice. For example, altered pitches cause spectral shifts (*bottom row panels*; **Fig. 8D**) that could scramble the feature assignments making them no longer effective. Hence, responses of the channel-selection method to the female voice show a peak that is nominally driven by the male voice, yet the model assigns it to the female output given the original label of these channels as female. Temporal coherence-based selection by contrast is not locked to a specific feature but instead primarily relies on the response *temporal* structure which remains a robust cue.

## Discussion

This study explored the neural mechanisms that underly the segregation of speech mixtures in ferret auditory and frontal cortex. Ferrets were trained to segregate speech mixtures consisting of word sequences uttered by simultaneous female and male voices. The ferrets either listened passively or attended to the female-voice and detected a target-word while ignoring the distractor male-speech. Two ferrets (**S, A**) learned to perform this task reliably while neural responses were recorded from primary (A1) and secondary (PEG) auditory cortices, and in the frontal cortex (FC) in ferret **S** and a third animal **C** who performed only a female-alone detection task.

### Speech segregation responses in the passive animal

When the animal was passively listening to the two voices, they were equally well represented in the auditory cortex, with cortical responses largely phase-locked to the syllabic modulations of the speech. However, single neurons tended to be more responsive to one voice or another depending on the match between the stimulus and the spectro-temporal selectivity of the cell. Thus, comparable numbers of cells in A1 and PEG could be labeled as female- *vs* male-cells, a clustering that simply reflected the difference between the spectro-temporal structure of the two stimuli in the voice mixtures. This distinction however was important because when the animals engaged in the segregation task attending to the female voice and detecting her target-word, there were consistent patterns of rapid plasticity in each cell group: female-cells became enhanced relative to the male-cell responses, either by boosting the female-cells responses or by suppressing the male- cell responses, or both. The net effect was to make the auditory cortex more responsive to the female-voice during task performance, and hence segregate it away from speech mixture.

### Speech segregation in human vs ferret auditory cortex

The findings above are consistent with previous ECoG studies that explored the segregation of 2-speaker mixtures in human auditory cortex [4,17]. For example, neurons in ferret auditory cortex exhibit a rich diversity of cortical STRFs just as in humans’ Heschel Gyrus (HG). Thus, if one cluster of cells is more responsive to one speaker or another, one can readily reconstitute the spectrograms of that speaker. Of course, this is not normally feasible because the responses to any one speaker are widely scattered and intermingled across the cortex, and hence can only be identified and grouped together based on other aspects of the responses such as the temporal correlations among them as we elaborate later.

A key difference between human and ferret cortical responses is their rapid plasticity once engaged in the segregation task. In humans, attention to one speaker or another apparently does not significantly affect HG responses [17], but only those of the secondary areas of the Superior Temporal Gyrus (STG) [17,4]. In ferrets, A1 responses by contrast become significantly more representative of the attended speaker, partly by relatively enhancing the responses of the cells selective to it or suppressing those selective to the distractor. These same effects but stronger are also seen in the ferret secondary auditory field (PEG). So unlike in humans, the response modulating effects of attention seem to be well articulated in ferret A1 and become even more pronounced in PEG (and presumably in FC) during task engagement.

### Speech segregation in relation to streaming

Speech segregation shares many of the perceptual characteristics typical of streaming tasks that have been extensively studied [1]. For instance, simultaneous sequences of simple tones, noise-bursts, or tone-complexes segregate perceptually (or stream apart) the more they differ in frequency, timing, loudness, rates, timbre, pitch, and bandwidths. Perceptual segregation is also commensurate with the listener’s ability to focus on one stream while ignoring the other. However, the most important determinant factor of streaming is the relative timing of the sequences in each stream. Thus, the less coincident they are, the more likely they are to stream apart perceptually [1,13].

A key feature of the streaming percept is its gradual buildup, often requiring a fraction of a second up to a few seconds to emerge. In our experiments, the ferrets needed to accomplish two perceptual goals following the onset of the speech mixture in each trial. First, to stream apart the speech mixture as quickly as possible so as to attend to the segregated female-stream. Second, it had to wait for the female-target word. This means that the animals had to suppress the male distractor throughout the trial in favor of the female-voice, both her reference and target words. This is indeed exactly why the female-reference words exhibited the same strong suppression of the male-voice both during the buildup of the reference sequence (**Fig.6 B, C)** and during the female-target word **(Fig.6D).** The details of the reconstructed spectrograms of **Figs. 4** provide a consistent alternate view of this plasticity.

### Streaming, segregation, and temporal-coherence

It is evident that cortical responses change during speech segregation in favor of the attended speaker. However, a difficult question arises: How does attention select the female- or male-cells for such opposite effects if the overall average response rates in the two clusters are initially comparable, and if the female- and male-cells are scattered irregularly all over the auditory cortex. How can the brain “know” which cells belong to one voice or the other, and relatively enhance the neurons that belong to the target speaker and/or suppress the others? A possible explanation is offered in the framework of the *Temporal Coherence* principle [12–14]. Briefly, the idea is that speech mixtures originating from independent voices are usually composed of different and persistently (*over 100’s milliseconds*) asynchronized sequences of words and syllables. For example, our male and female syllables differed in pitch (spectral harmonics) and timbre (formants). But crucially, they differed in their timing, with dissimilar or alternating onsets between the male and female sequences (**Fig. 1**). Therefore, cells tuned to the different spectral features of the male or female syllables also exhibited different temporally modulated responses. Hence, a cluster of male-driven or female-driven cells tended to have co-modulated responses within the group but were uncorrelated or negatively correlated with the responses of the other group. This explains how it was possible to cluster the responses in A1 and PEG into two groups based on their correlation patterns (**Figs. 5**). It is this segregation of correlated responses that allowed us to reconstruct and observe the contributions of the two speakers to the different frequency channels and the diverse effects of attention on them (**Figs. 4**).

The other critical ingredient for segregation to occur is the notion of “binding”, namely that highly correlated cells (within the female- or male-cluster) rapidly form excitatory inter-connections that mutually enhance their responses. The opposite happens between anti- or weakly-correlated cells which form mutually inhibitory connections that suppress the cells [27, 38]. Thus, in the case of a mixture of two speakers, if a listener attends to one voice, e.g., the ferret attending to the female voice, all it needs to do is boost the responses to one distinctive attribute, e.g., the pitch or location (if they are separated). This in turn would enhance all other responses that are correlated with those of the female voice, e.g., responses to the harmonic components in the stimuli [28]. The enhanced female responses suppress the competing uncorrelated responses of the male voice. These changes are consistent with many details of our findings. Furthermore, we find that the onset of rapid plasticity occurs only when the animal engages in the task and attends to the female voice, thus inducing sustained effects throughout the duration of the speech stream and causing simultaneous enhancement of all female words, and suppression of all male words in the mixture (**Figs. 6**).

Overall, our experimental findings and their computational modeling suggest that temporal coherence is a fundamental process to explain how the auditory system can segregate and attend to a desired target sound in a mixture. At its early cortical stages, the auditory system relies on richly diverse feature selectivity (e.g. frequency, timbre, location, pitch) [34, 37, 39] to disentangle and represent sound mixtures along different attributes. In the case of concurrent speakers, each speaker would thus dominate the responses of a set of matched cells in A1, and with correlated temporal response patterns that reflect the syllabic sequences of that speaker’s words. To tie these responses together as those belonging to a common source one wishes to listen to, *temporal coherence* postulates that correlated neural activity promotes excitatory connectivity (cooperative) and mutual enhancement of the responses among these neurons, while suppressing uncorrelated responses through inhibitory (competitive) interactions [38]. Ultimately, the attended voice emerges through this enhanced alignment [34], consistent with rapid plasticity occurring within fractions of a second and commensurate with typical perceptual buildup of streaming [17, 36, 38] and also the sustained effects throughout the duration of the speech stream causing simultaneous enhancement of all female words, and suppression of all male words in the mixture. Of course, such a temporal coherence hypothesis needs more direct support from *in-vitro* measurements of neuronal connectivity while neurons are driven coherently or incoherently.

## Supplementary Figures

**Supplementary Figure S1.**
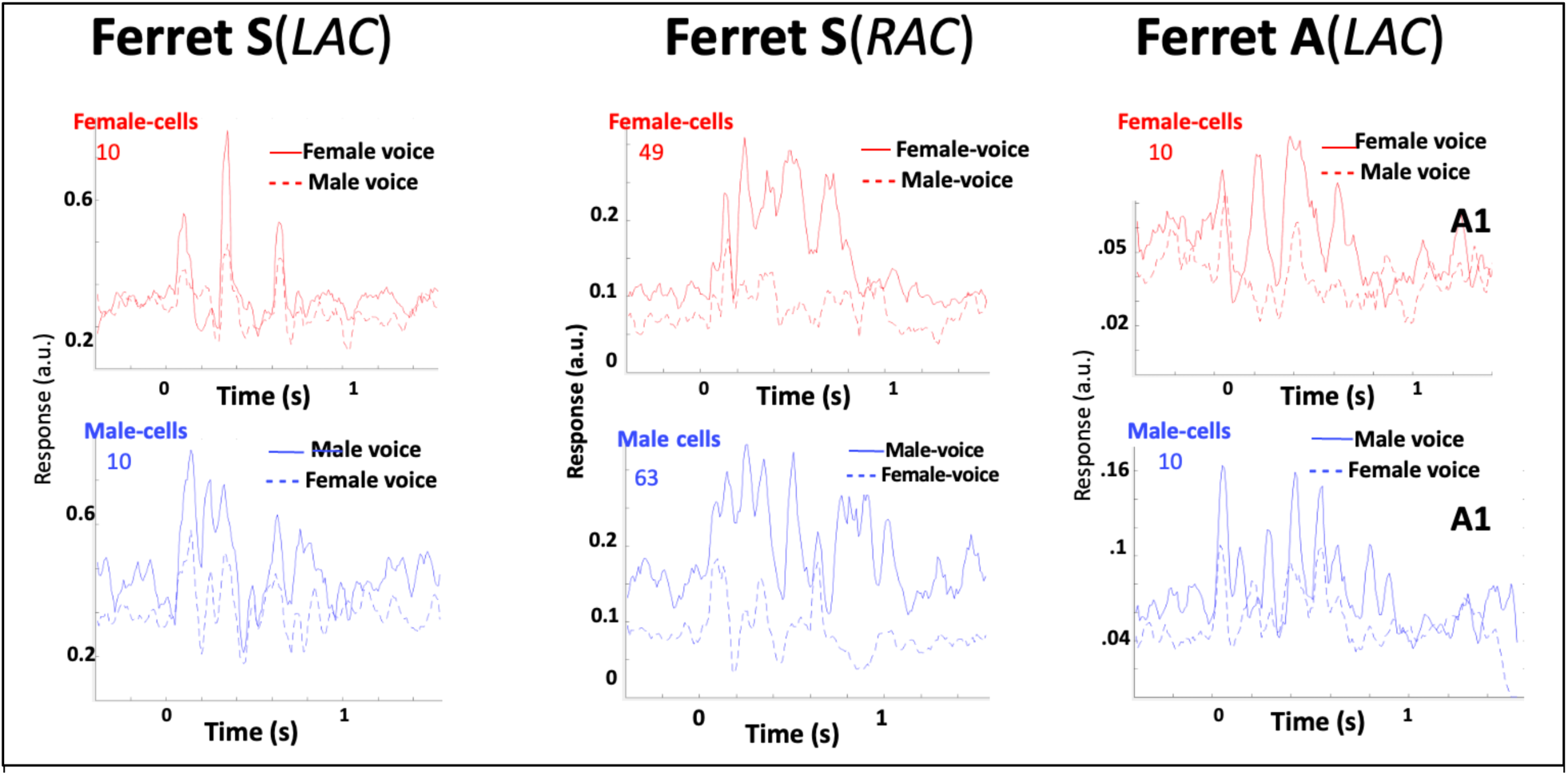
Examples of PSTHs from a selection of cells from each of the 3 hemisphere of ferrets S & A. All cells are characterized by their *SSI* and grouped into female- and male-cell clusters. As expected, they demonstrate that the responses of a cluster is stronger to the male/female voices that define it. Note that the words used are different in the different experiments and hence it is not always possible to meaningfully combine the responses from all cells into one PSTH.

**Supplementary Figure S2.**
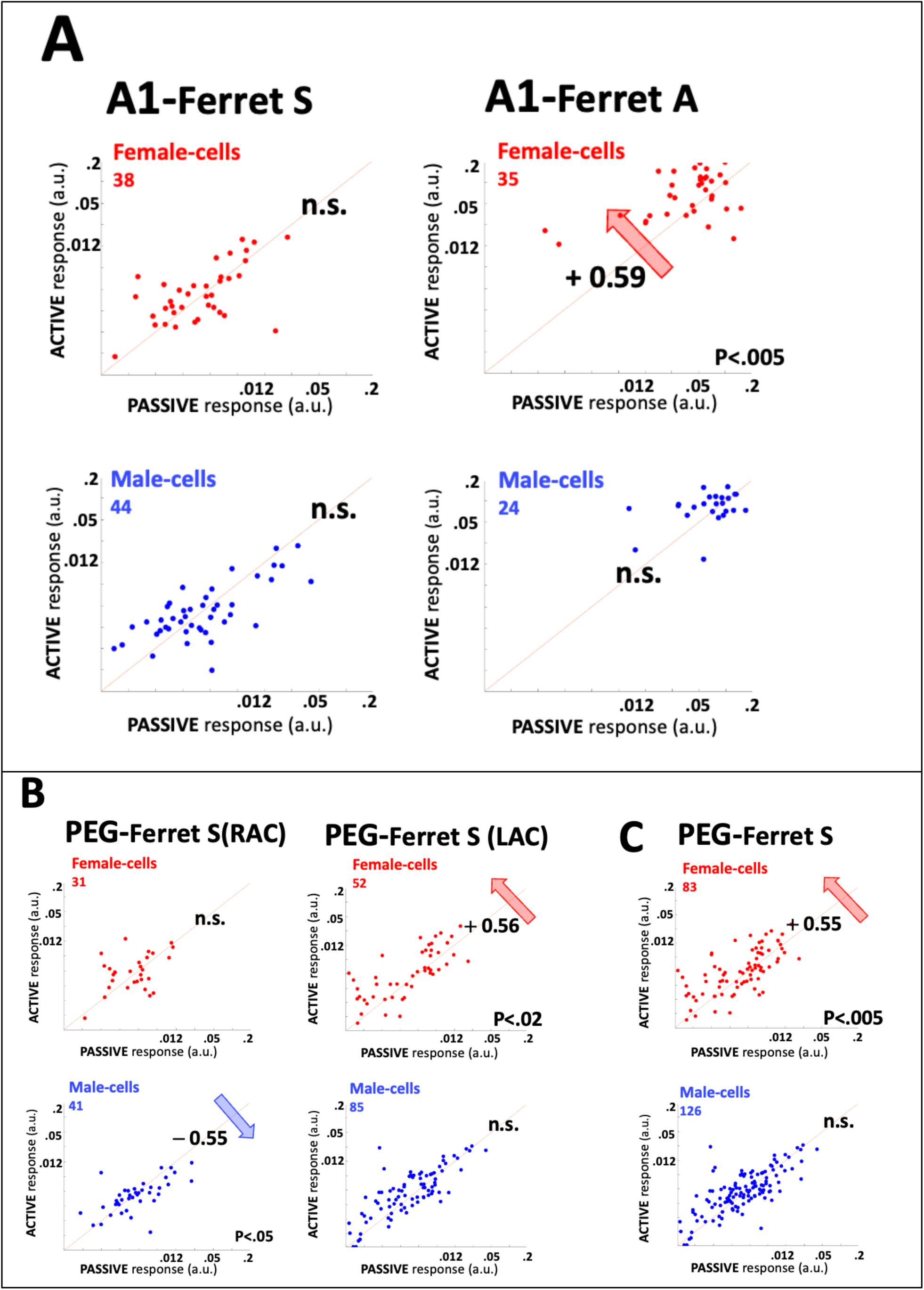
Distribution of plasticity in male- and female-cells across the A1 and PEG in auditory-hemispheres in 2 ferrets: **(A) A1 plasticity in 2 animals.** It exhibits female-cells enhancement in one animal. Male-cell plasticity is absent. **(B) PEG plasticity in 2 hemispheres of ferret S.** In one hemisphere (RAC) only male-cell suppression is seen. In the other (LAC), only female-cell enhancement is present. **(C) PEG plasticity in PEG of ferret S.** The combined plasticity of all PEG cells here is significant only in female-cell enhancement.

**Supplementary Figure S3.**
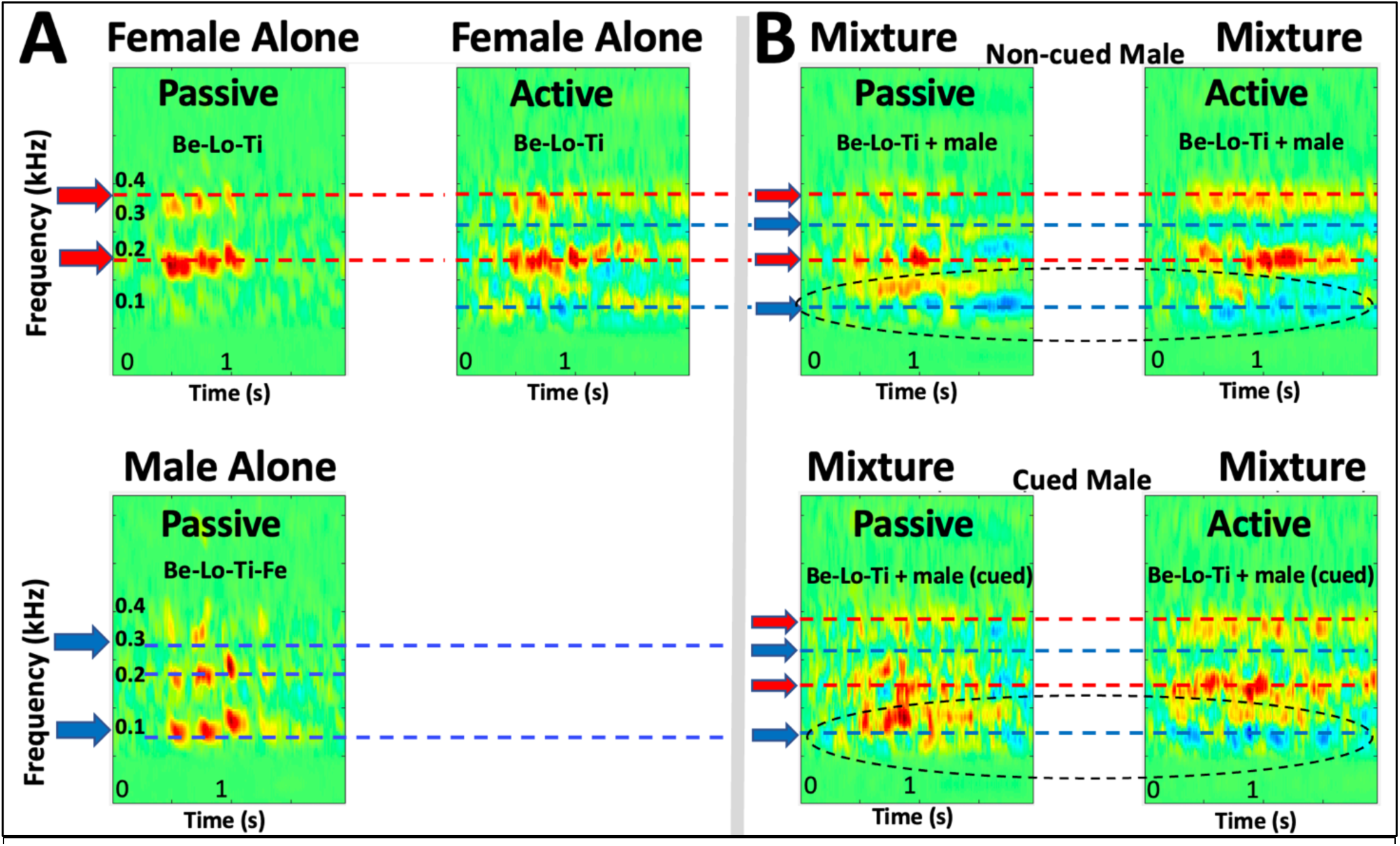
Reconstructions of the PEG responses in ferret A. **(A) Reconstructions of single voice responses.** The reconstructions in the passive state resemble the spectrograms of the stimuli, namely the male and female target-words. When the animal is engaged in detecting the female-alone target, the responses remain as in the passive, but with an additional overall surround suppression. **(B) Reconstructions from male/female mixtures.** During the passive state, the responses to the mixture combine the spectral features of both voices (*left panels*). When the animal is in the active state (*right panels*), the male- (distracter voice) channels are evidently suppressed compared to the passive state. In the bottom panels, the displays are aligned (cued) to the male-voice words. This requires slight temporal shifts from trial to trial since the male-words are (by design) misaligned with the female-words. The reconstructions here display the same prominent features described in the top panels, with suppression of the male responses during the active state.

**Supplementary Figure S4.**
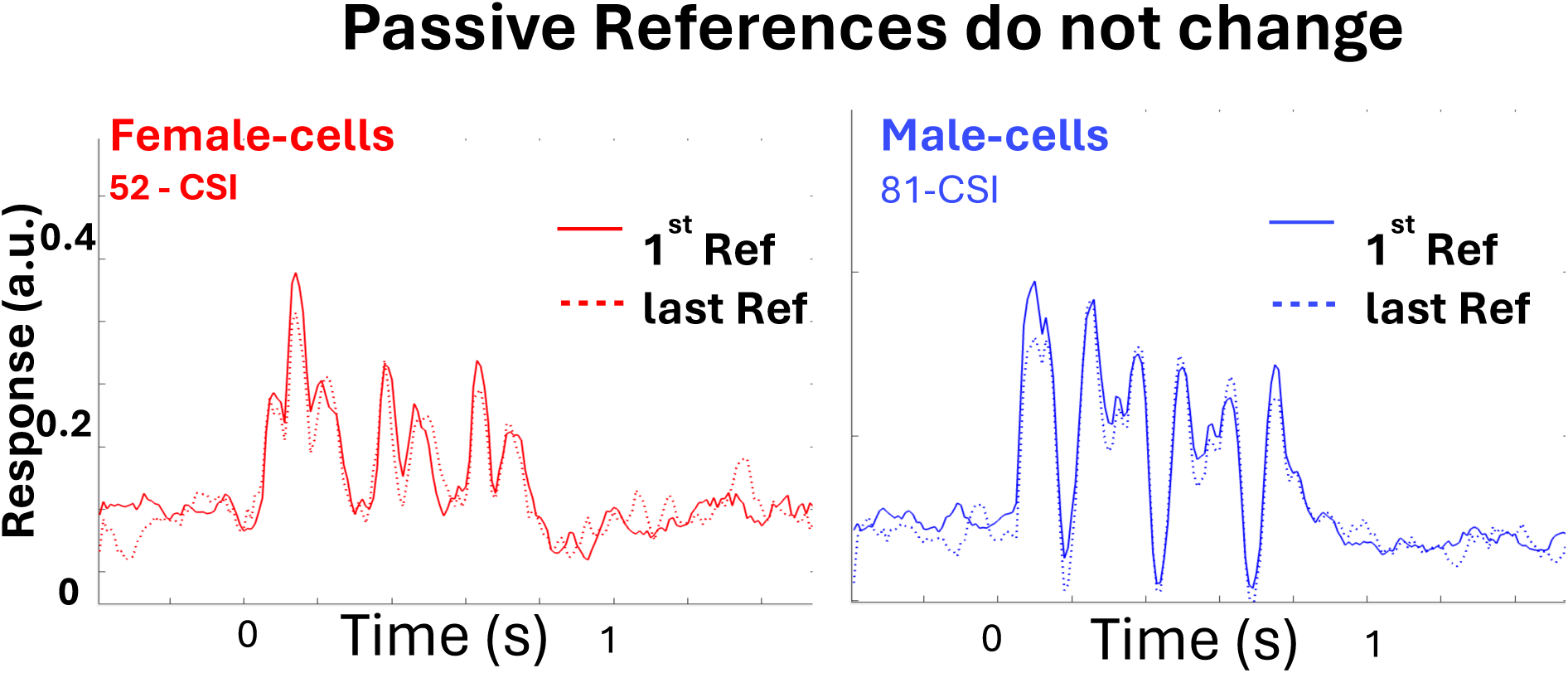
Reference sequence in ACX of ferret S: Reference responses in the passive state remain essentially unchangde throughout the sequence.

**Supplementary Figure S5.**
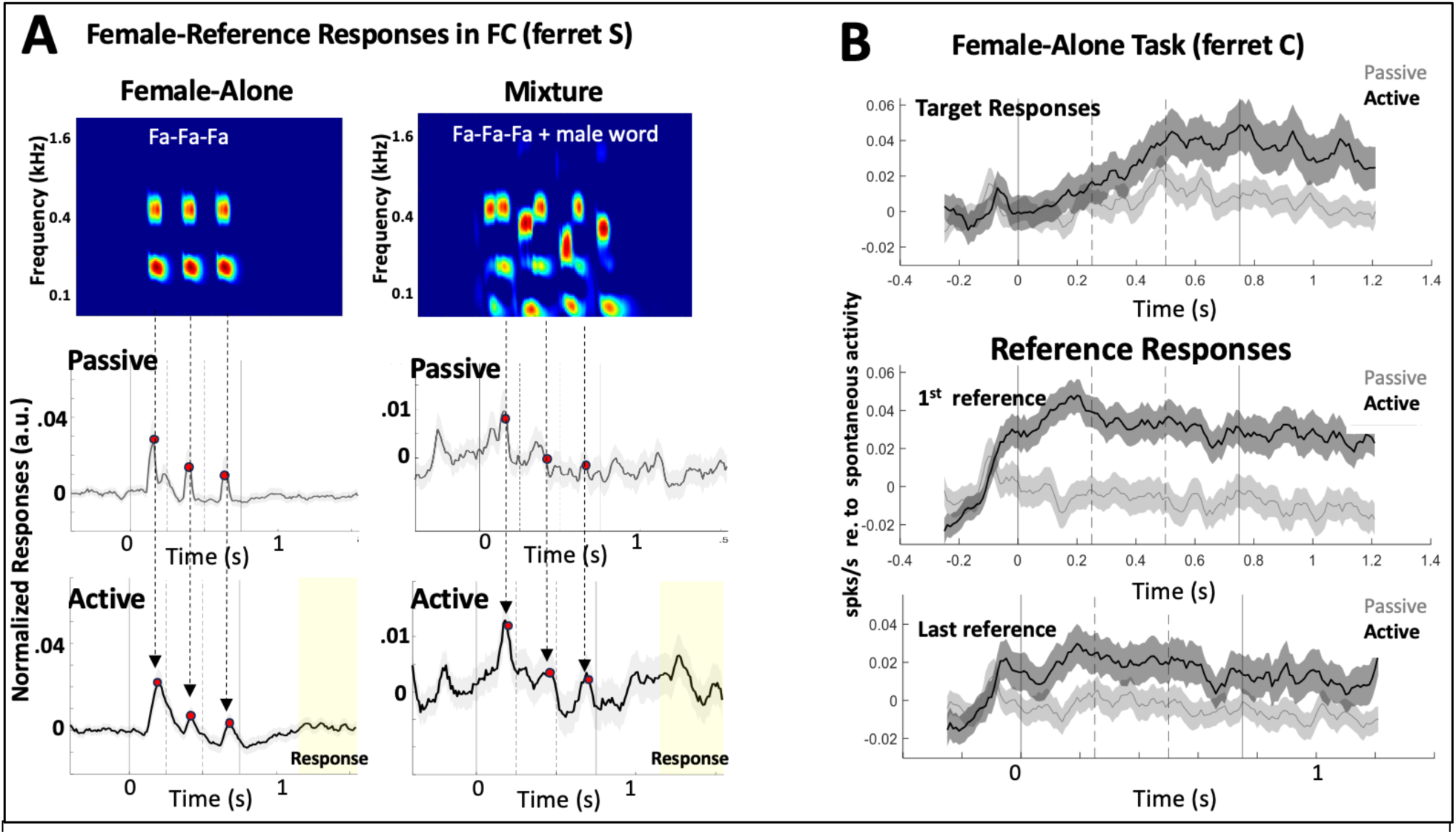
Female-reference word responses in FC. **(A) Female-alone and mixture tasks.** Syllabic response peaks of reference word /FA-FA-FA/ become enhanced relative to the baseline during task engagement. Overall, plasticity in reference-words recapitulates the findings from the target-word in that passive responses are weak (*left-middle panel*), becoming stronger during active performance (*left*-*bottom panel*). During the mixture segregation, the female-reference responses are cluttered by the interfering male-words (*right*-*middle panel*) becoming more visible during the active state (*right-bottom panel*) as the male-peaks are suppressed. **(B) Female-alone responses in FC of ferret C.** Enhancement of the target responses in FC is typical of other auditory cortical areas *(Top panel).* Reference responses in the FC gradually decrease from 1^st^ to last reference *only* during engagement (*middle and bottom panels)*. By contrast, in the passive state they remain unchanged (middle *versus* bottom panels).

**Supplementary Figure S6.**
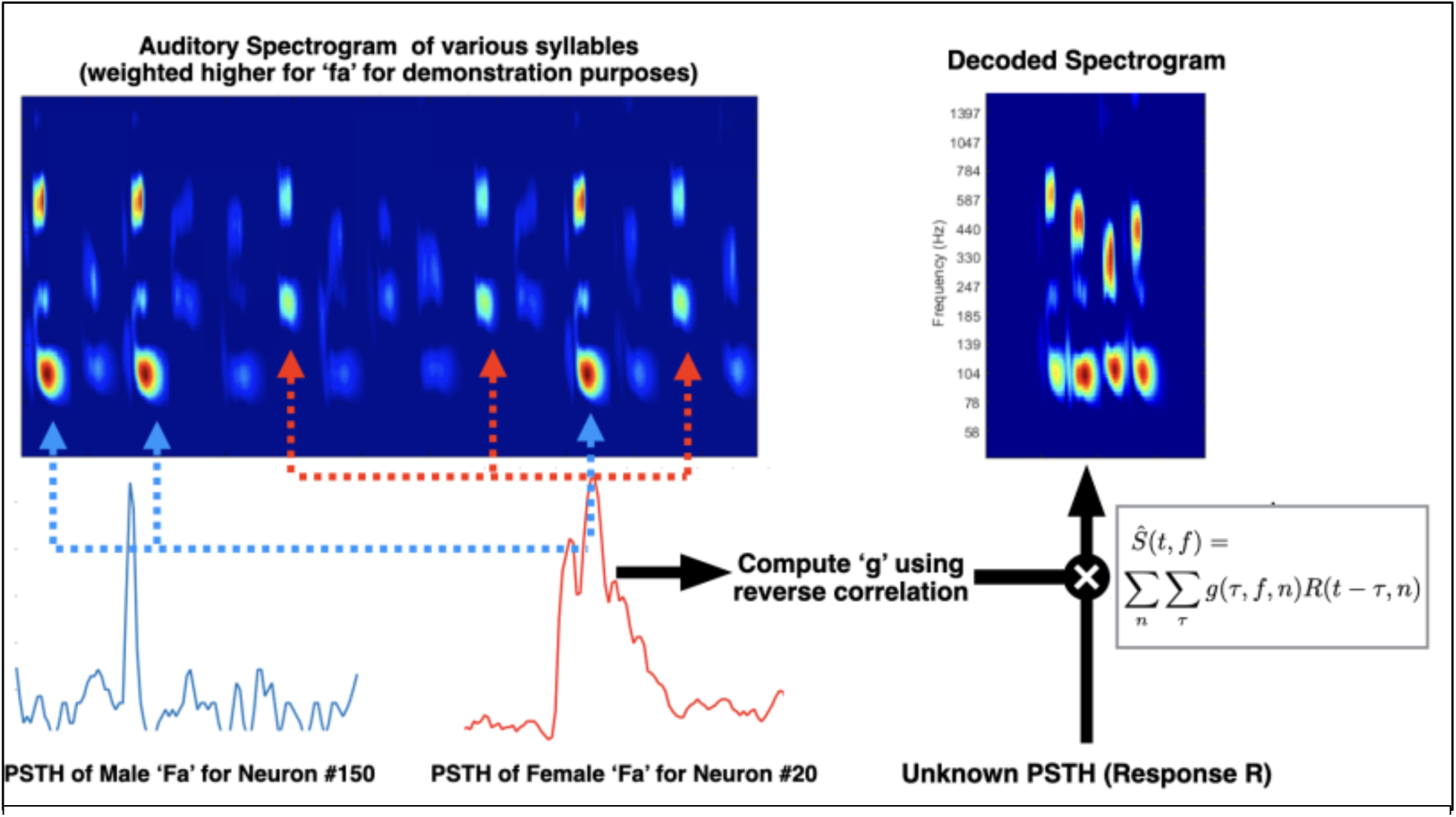
Stimulus reconstruction. Syllabic responses were collected per neuron, along with their auditory spectrograms. Linear filters were obtained through reverse correlation, to finally get the decoded spectrograms from PSTHs.

## MATERIALS AND METHODS

### Animals

Three adult female ferrets (Mustela putorius, Marshall Farms, North Rose, NY) were trained for the neurophysiological experiments, with two trained on both female-only and mixture conditions (ferret **A** and ferret **S**), and one trained on only female-alone condition (ferret **C**). The animals were placed on a water-control protocol in which they obtained water as rewards during behavior sessions or as liquid supplements if the animals did not drink sufficiently during behavior. They also received ad libitum water freely over weekends. Animals’ health was monitored, and they were maintained above 80% of their ad libitum weights. Ferrets were housed in pairs or trios in facilities accredited by the Association for Assessment and Accreditation of Laboratory Animal Care (AAALAC) and were maintained on a 12-hour light-dark artificial light cycle.

All animal experimental procedures were conducted in accordance with the National Institutes of Health’s Guide for the Care and Use of Laboratory Animals and were approved by the Institutional Animal Care and Use Committee (IACUC) of the University of Maryland.

## EXPERIMENTAL PROCEDURES

### Behavioral Tasks

The task in both training and neurophysiological experiments was a conditioned avoidance (Go/ NoGo) task in which the ferrets had learned to continuously lick the sprout when the reference words were presented (Go Phase), and to refrain from licking when the female target word was uttered (NoGo phase). The target word took the form ABC, where A, B, and C referred to three distinct syllables, whereas the female reference words could be AAA (ferret **S**), ABX (ferret **C**), or AXY/AQR (ferret **A**). In the female-alone condition, each typical trial consisted of the female trisyllabic references presented randomly between one to six times then followed by the female target word. About 15% of all trials were catch trials, where the references were uttered two to seven times with no target word in the trial. The words were separated by an interstimulus interval of 1.2 seconds.

The mixture condition trials were similar to those of the female-alone condition, except male quadrisyllabic words were presented simultaneously with the female stream. The animals were taught to pay attention to the female speaker and ignore the male speaker. For each female utterance, a male word randomly chosen among seven quadrisyllabic words would be presented simultaneously. One of the seven words explicitly contained the target sequence to allow control analyses of behavior to the unattended male speaker. To simulate the partial overlap and temporal incoherence of speakers typical in real-life situations, a relative delay between the background male stream and the foreground female stream was introduced. Each of the background male words were shifted by starting them either 400ms before, 80ms before, or 200ms after the female words.

In both the female-alone and mixture conditions, a 400-ms buffer window was provided for the animals to decide. If the animals licked in the reinforcement period following the target word (400ms – 800ms after word ended), they were presented with a mild shock on their tails (paws during training) to solidify the target word as a NoGo word.

Table 1 provides the female target, female reference words, and male words used for each animal. Note the set of male words were shared across animals that were trained on the mixture condition.

**Table 1.**
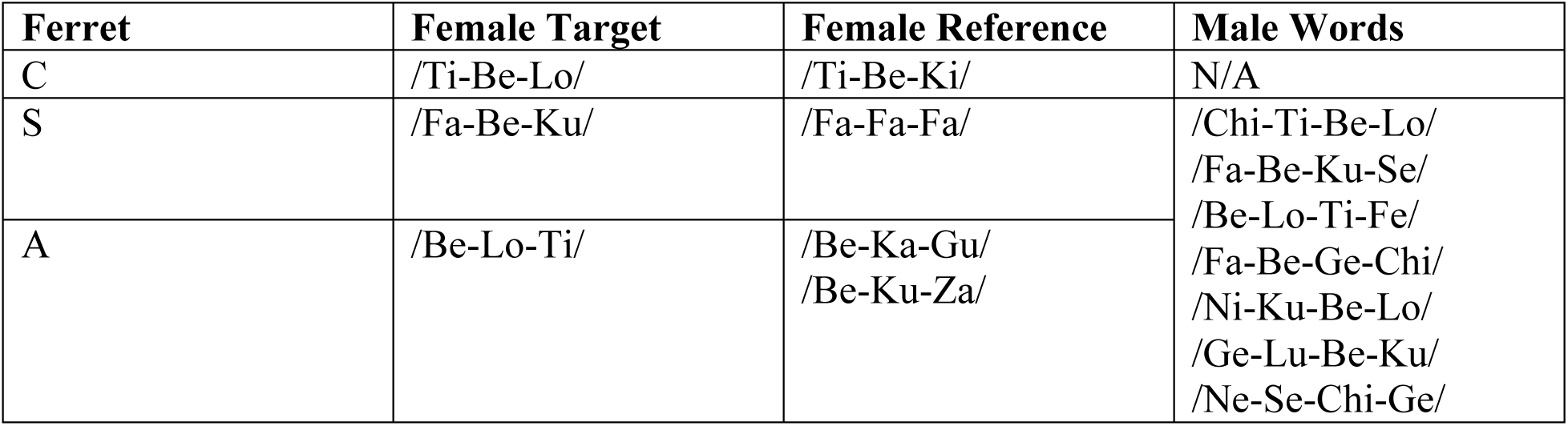
List of female target, female reference, and male word stimuli.

### Training

Animals were initially trained in a free-moving setup where they learned to lick from a sprout at the front of a Faraday cage. Water flowed continuously through the sprout at a rate between 0.3 mL/min to 4 mL/min. Training occurred with intermediate steps to confirm performance at an easier version of the task before progressing to the final version of the task design. Animals started learning the task with a single speaker, with the target word presented up to 30 dB louder than the reference words. As they learned the task, determined by the consistent accuracy of performance across three consecutive training sessions, the second (male) speaker was introduced at lower loudness. The loudness of the second speaker was increased as the animal learned the task until the performance was consistent when the two speakers were equally salient. Once they performed consistently above chance level at the final version of the task design, where consistency was defined as Accuracy (Hit Rate) = 75%, Consistent Licking pre-stimuli (Safe Rate) = 50%, and Discrimination Rate (Hit rate controlled for inconsistent licking) = 40%, we considered the animal ready for implantation. More specifically,

i. Hit Rate (HR) = Number of trials of successfully refraining licking at Target divided by the Total number of trails with Target. The chance level of HR is 0.5.
ii. Lick Rate or Safe Rate (SR) = Amount of time the animal licked before word presentation. This is used to determine whether the animal actually stopped licking due to the Target, or if the animal had a NoGo trial by the virtue of not licking during and prior to the Target presentation. The latter case would entail no actual Target detection was involved, and such trials were labeled as “snooze” trials whose responses were not counted towards successful NoGo trials. The chance level of SR is 0.5.
iii. Discrimination Rate (DR) = Accuracy modified by Lick Rate, given by DR = HR * SR. Therefore, the chance level of DR is 0.5 * 0.5 = 0.25.

### Headpost Implant Surgeries and Neurophysiological recordings

After reaching behavioral criteria on the task, a stainless steel headpost was surgically implanted on the ferret skull under aseptic conditions while the animals were deeply anesthetized with 1% - 2% isof1orane. The headpost was secured in the skull using titanium screws and embedded in Charisma (Ferrets S and C) or dental cement with Gentamicin (Ferret A). The area around the auditory and frontal cortices were covered in a single layer of cement, whereas surrounding areas were covered with 5 – 7 mm-thick cement to protect the exposed skull.

After recovery from surgery (about 2 – 3 weeks), the animals were placed in a double-wall soundproof booth (IAC) and were habituated to the head-fixed setup. They were re-trained in the head-constrained version of the task, where a mild shock was delivered to the tail on “miss” trials. Neurophysiological recordings began after the animals regained consistent criterion levels of performance (Hit Rate = 75%, Safe Rate = 50%, Discrimination Rate = 40%) in three sequential behavioral sessions. All behavioral data in this paper was obtained after implantation, including regaining performance to pre-surgical levels.

To expose a part of the auditory or frontal cortex for recording, small 1 – 2 mm craniotomies were made in the skull. Recordings were conducted using 4 tungsten microelectrodes (2-5 Momaga, FHC) simultaneously advanced through the craniotomy and controlled by independently movable drives (Electrode Positioning System, Alpha-Omega) with 1 micrometer precision until well isolated spiking activity was observed. Raw neural activity traces were amplified, filtered, and digitally acquired by a data acquisition system (AlphaLab, Alpha-Omega). Single units were isolated offline using customized spike-sorting software based on PCA, K-means clustering, and subsequent template matching. Only units with greater than 70% isolation and a typical refractory period were conserved as single units.

### Experimental Procedures and Stimuli

Each recording began with presenting 250-ms random tone pips of varying frequency (125 – 32000 Hz, 4 tones/octave) and intensity (0 to -50 dB range, 10 dB increment) to determine the characteristic frequency (CF) and latency of each individual recording site. A typical recording would then include two passive female-alone blocks that sandwiched the female-alone behavioral block (Ferrets S and A, repeated twice for Ferret C), and two passive mixture blocks that sandwiched the mixture behavioral block (Ferrets S and A).

Speech was generated using the Straight synthesizer for speech for Ferrets S and C, and the inbuilt Apple synthesize for Ferret A. The fundamental frequency (F0) of the male speaker is 107 Hz, and that of the female speaker is 220 Hz for all stimuli presented to Ferrets S and C, and the F0 of the male speaker and the female speak is 100 Hz and 180 Hz respectively for Ferret A. Each syllable is an English syllable of the form “CV”, comprising of a consonant followed by a vowel, with a length of 180 – 220 ms. All syllables are then made to be of a constant length of 250 ms to control for effect of syllable duration response before creating words. Words are comprised of three to four of these syllables concatenated with a 10-ms onset and offset ramp to avoid transient transitions. Since the animals were trained to recognize the female target word, the trisyllable words were uttered by female and quadrisyllabic words were uttered by male. The same set of stimuli was used across recordings in A1, PEG, and FC across the passive and behavioral conditions within an animal.

The spectral resolution of the spectrograms as shown in Fig.1A, and used for the analysis and reconstructions explained below, was set to reflect the moderate frequency tuning of the ferret auditory system, typically high-enough to resolve the lowest 2-3 harmonics [20–22]. Consequently, spectrograms clearly depicted the resolved fundamental and up to the 3^rd^ harmonic, as well as the syllable formants (**Fig. 1A**). All syllables consisted of a consonant-vowel (CV) combination with voiced segments of different pitches with several harmonics of fundamental frequencies 100/107 Hz (male) and 180/220 Hz (female) in ferrets **A** & **S** experiments, respectively. These choices ensured that female and male voices dominated different frequency channels (male dominated ∼100 and 300 Hz, or 1^st^ and 3^rd^ harmonics; female dominated ∼200 and 400 Hz, 1^st^ and 2^nd^ harmonics). This segregation facilitated the visualization and assessment of enhancements and suppression of the neural activity associated with the two competing voices.

All acoustic stimuli were presented at 65 SPL except for the tone pips. The sounds were digitally generated using custom-made MATLAB functions at 40 kHz sampling rate and were converted at 16-bit resolution through a NI-DAQ card, then amplified and delivered through a free-field loudspeaker located ∼1 m in front of the animals’ head.

### Localization of recording sites/ Identification of recording sites by tonotopic

Recording locations in each ACX hemisphere were characterized by their locations along the dorso-ventral and rostro-caudal axes. Artificial marks were created by drilling a depression in the headcap on either side of the craniotomy as reference landmarks for localizing recording positions. Furthermore, the neuron’s CF was measured at each electrode penetration. The CFs obtained from all penetrations were then aligned to form a tonotopic map for each animal, based on which the locations of A1 and PEG were confirmed. The CF gradient in A1 runs from high to low frequency along the dorsal-lateral direction, and the gradient direction was reversed at the low frequency border with PEG. Therefore, the lowest CF contour line was used as the dividing border between A1 and PEG. Neurophysiological data was also recorded from FC. Since FC does not demonstrate evidence for tonotopy by frequency, the locations of recordings were measured by referencing the two landmarks placed in the bone cement surrounding the craniotomy.

## DATA ANALYSIS

### Computing characteristic frequency (CF)

The characteristic frequency (CF) of each neuronal unit was obtained by analyzing their responses to tone pips with varying frequency and intensity (see Experimental Procedures and Stimuli). A two-dimensional frequency x intensity response matrix was then created by taking the mean evoked response to tones at each frequency and intensity level. The response matrix was baseline-corrected by subtracting the mean and dividing by the standard deviation of the baseline activity from 100 ms before tone onset. The longest iso-response contour line in the normalized matrix was defined as the neuron’s tuning curve, and the frequency corresponding to the lowest intensity on the tuning curve was the neuron’s CF. A penetration site’s CF was computed by taking the median value of all isolated single units in that site.

### PSTH calculations

Peri-stimulus time histogram responses (PSTHs) were obtained by binning the neural responses into 10-ms time bins and averaging these windowed spike data across trials and stimulus presentations per cell, then further averaging across cell. All shaded error bars represented the standard error of mean (s.e.m.) on the cell level. Unless otherwise specified, responses to target only included hit trials, i.e., trials where the animal successfully refrained licking upon detecting the target words, whereas responses to reference included all trials.

Unless otherwise specified, PSTH responses used in further analysis are without baseline correction nor normalization. PSTHs were plotted without baseline removal and normalization, unless specified as “referenced to spontaneous activity,” in which case the average of neural responses during a 250-ms pre-stimulus time window was subtracted from the PSTH for each cell before averaging over population.

### Speaker Selectivity Index (SSI)

The Speaker Selectivity Index (SSI) is a device that quantifies the coincidence between a cell’s BF and the spectral components of the stimuli. The SSI for each cell is defined as the normalized difference between the variance of the cell’s average male-alone response and the variance of the cell’s average female-alone target response, both in the passive condition. A value of -1 < SSI < 0 represents the female-voice preferring cells, and 0 < SSI < I for the male-voice preferring cells.

### Correlation Selectivity Index (CSI)

ACX neurons are usually able to phase-lock to the modulations of their most preferred stimulus. Thus, the responses to the female-preferred and the male-preferred clusters can also be differentiated by their mutual correlation, that is, responses of the female-cells should be more correlated within themselves than with male-cells, and vice versa. To demonstrate such segregation, the Correlation Selectivity Index (CSI) was calculated to obtain the strength and sign of each cell’s relative contribution to the overall phase-locked response. The CSI was computed by finding the covariance between the average passive responses of all pairs of ACX cells to the mixture of female and male words and subjecting the covariance matrix to singular value decomposition. The weights of the largest eigenvector were used to cluster the cells, such that cells with *positive* eigenvalues were assigned to the female cluster and cells with *negative* eigenvalues were assigned to the male cluster.

### Response Power

The response power of a cell in the passive or behavioral condition was quantified as the variance of the mean response of the cell to the *mixture* of female and male words under said condition. To capture the effect of task engagement on population plasticity that is reflected through the strength of temporal modulation, we measured the *effect-size* of a population, by first plotting the response power of each cell on a scatter plot with the passive response power on the x-axis and its active counterpart on the y-axis. In such a plot, the diagonal line with a slope of 1 would represent no effect of task engagement since the variance is consistent across conditions. A cell lying under the diagonal line would mean the amplitude of the temporal modulation of that cell is diminished under the behavioral state, and above the diagonal line would mean the active state is correlated to an increase in the strength of temporal modulation. To assess the population-level effect, the *effect-size* was computed by finding the signed distance of each point on the scatter plot from the midline, normalizing all signed distances by the mean of the unsigned-distances, and averaging all normalized signed distances in the population. The *effect-size* therefore takes a range of -1 to +1, with negative values representing an overall suppression of the active responses across cell population, and positive values indicating an overall enhancement of responses in the active state relative to the passive state.

To determine if a scatter-asymmetry is significant, a one-sample t-test of the distribution of all points in a scatterplot around a midline is computed to determine if its mean is shifted significantly above or below the midline. The t-test (MATLAB *ttest* command) is evaluated on the distribution of normalized signed distances. For comparison between the *effect-sizes* of two conditions (e.g., active versus passive), a two-sample t-test (MATLAB *ttest2* command) is applied on the two distributions of normalized signed distances to see if they are significantly different. Significance in either test is defined by a p-value < 0.05.

### Stimulus Reconstruction

The population PSTH responses and effect-size revealed strong indications that changes in plasticity reflected in the strength of temporal modulation may contribute to the segregation of speech mixtures and the enhanced perception of the attended female stream. To understand their contributions to the details of the speech segregation in both the time and frequency domains, the method of linear reconstruction of the stimulus [23] was employed. Responses to individual syllables used to construct the female and male words were first obtained for each neuronal unit. We then trained a linear *inverse* filter that mapped the responses to the stimuli by computing for each cell the correlations between the obtained responses and the stimuli’s corresponding auditory spectrograms while minimizing the squared error between the reconstructed and stimulus spectrograms. The auditory spectrogram is a time-frequency representation that models the early auditory processing stages of the auditory pathway, as outlined in [29]. We then reconstructed the spectrograms implied by the responses while the animals listened to the mixture passively or during active engagement in segregating the speech mixture. The resulting reconstructed spectrograms thus represented the encoded neural responses of the population, and comparing the two spectrograms from passive and active conditions would reveal the effect of task engagement and how the ACX extracted the attended female-voice.

More specifically, the original sound stimulus was represented in auditory spectrogram 𝑆(𝑡, 𝑓), where 𝑡 = 1, …, 𝑇, T being the length of the signals, and 𝑓 = 1, …, 128 channels spanning from 50 Hz to 4000 Hz. For a population of 𝑁 neuronal units, we represented the response to stimulus of neuron 𝑛 (1 ≤ 𝑛 ≤ 𝑁) at time 𝑡 = 1, …, 𝑇 as 𝑅(𝑡, 𝑛). The neural responses had been processed to remove baseline firing activity. The *inverse* function 𝑔(𝑡, 𝑓, 𝑛) is a function mapping the response 𝑅(𝑡, 𝑛) to stimulus 𝑆(𝑡, 𝑓) as follows:

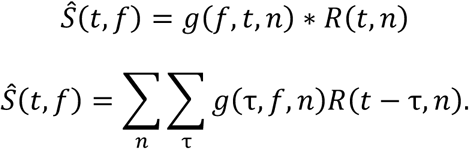

Since the reconstruction at a particular frequency 𝑓 is independent of the other frequency channels, the following analysis would focus on a single frequency channel 𝑓, and the exact analysis could then be replicated with the rest of the channels. Let *g_f_* be the channel *inverse filter*, and it is estimated by minimizing the mean-squared error *e_f_* between the original stimulus auditory spectrogram *S_f_* (*t*) and the reconstructed spectrogram 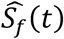 for that particular frequency channel, namely

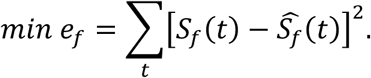

Solving the minimization problem yields a closed form solution involving normalized inverse correlation, given by

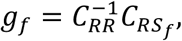

where

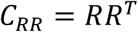

and

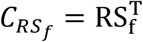

are respectively the auto-correlation of neural responses and the cross-correlation between neural responses and stimulus at different lags. *R* and *S_f_* are defined as

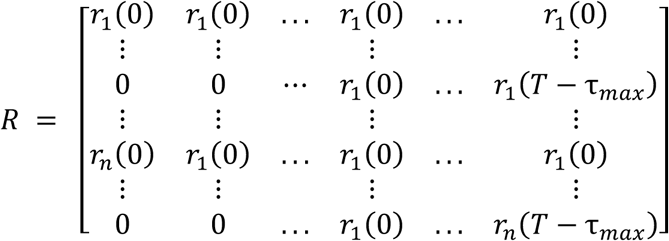

And

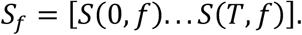

The matrix 𝑅 is padded with zeros on the left to insure causality. The matrix *C_RR_* is full rank due to the stochastic and causal nature of the neural response, hence is invertible. We experimented with different sets of causal and non-causal lags, with which no difference in the results for our tasks was observed. Lags from 0 to *τ_max_* = 200 ms in steps of 10 ms were chosen at the end. The entire reconstruction filter is then composed of functions of each spectral channel to obtain

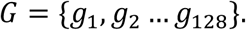

### Computational Modeling

A framework for source separation was adopted for the current study [34]. The model was previously shown to work effectively for steam segregation and was compared to state-of-the-art models for both speech and music segregation. It was adapted in the present study to align with speech materials used for ferret experiments. Specifically, the model takes as input the log short-term Fourier Transform (STFT) magnitude of the sound mixture (sampled at 16 kHz) and an indicator variable denoting the intended stream of interest. STFT is calculated using a window size of 4098 and a hop size of 512 with a hamming window (40 frames). The indicator is a unitary vector representing which memory to select for further processing of speech. The model was trained to learn three memory states corresponding to Male attention, Female attention, and No attention. In the first two cases, the model outputs a spectrogram emphasizing the target voice. In the third, the model operates as an autoencoder and outputs the mixture back. The first part of the model is a feature analysis stage which consists of a convolutional layer, a max-pooling layer, and a dilated convolutional stack followed by an up-sampling block. All max-pooling operations were performed using a kernel size 2 and stride 2. Each convolution node consists of 128 hidden units, kernel, and “same” padding with leaky *ReLU* (rectified linear unit) activation. A learning rate of 0.001 was used to train the model for 10K steps with a batch size of 64 on am NVDIA V100 GPU.

The next stage of the model consists of a coherence-based attentional selection which acts as a form of filter to modify the model embeddings in line with the attentional goal of the model. This gating operation uses target anchors (learnt during the training phase) onto which all embeddings are projected. During training, the model identifies defining characteristics of a specific target class (e.g., male voices) and associates relevant coherent auditory features with this anchor over time. These anchors essentially act as the attentional focus. They are not simple channel labels as specific targets do not always occupy the same channels (e.g. with shifted spectra). This memory of specific targets (spanning both frequency and embedding dimensions) is then deployed during testing when an unseen mixture is given as input. The role of this memory is to anchor activity from the incoming mixture such that neurons that are phase-aligned with this anchor will be enhanced while those that are uncorrelated over time are suppressed.

#### Training Data

The model was trained using the speech materials used for ferret experiments, specifically ferret **S**. The fundamental frequency (F0) of the male speaker is 107 Hz, and that of the female speaker is 220 Hz for all stimuli presented to the model. Each syllable is an English syllable of the form “CV”, comprising of a consonant followed by a vowel, with a length of 180 – 220ms. All syllables were aligned to a constant length of 250ms to control for effect of syllable duration response before creating words. Words were comprised of three to four of these syllables concatenated with a 10-ms onset and offset ramp. As in the ferret speech material, the female contained 3 syllables and the male contained 4 syllables The mixtures were created with a delay chosen at random from -400ms, -80ms and +200ms relative to the onsets of the female words. An independent (non-overlapping) subset was generated to evaluate the model performance.

#### Quantifying effects attentional gating

First, model neurons were ‘labeled’ as male or female neurons by analyzing their responses (pre-attention) to single voices using the same definition of speaker-selectivity index (SSI) defined earlier. A different labeling was also considered using the response of pre-attention neurons to the input mixture based on their inter-neuron correlations, following the same definition of correlation selectivity index (CSI) defined earlier. Since the model has access to >10,000 neurons overall, all analyses were performed with the top 500, 400, 300, etc. neurons based on SSI (or CSI) selection for male versus female neurons. Statistical power was evaluated for different number of neurons selected and results were overall qualitatively consistent regardless of number of neurons selected, though statistical power starts dropping below significance when number of neurons drops below 50. Results reported here use 200 neurons.

Once labeled (through the SSI or CSI indices), each cell cluster (male or female) was then analyzed pre- and post-attention to compare changes in response characteristics for a given attentional state (male or female attention). We quantified mutual information (MI) between responses of a given neuron across time and different stimulus trials (different input mixture delays and speech syllables), as a measure of mutual dependence between responses before and after attention. Mutual information was implemented using two different methods which yielded qualitatively similar results: using binning for the estimation of the probability distributions or using nearest neighbor estimations following the procedure in [37]. In both methods, we used cross-validation to confirm choice of estimation parameters. *Effect-size* was then quantified using the same process defined earlier to quantify influence of male versus female attention on male and female neurons separately. *Effect-size* quantified the signed distance of each point from the midline, normalized by the mean of unsigned distance. A one sample t-test (MATLAB *ttest* command) evaluated the distribution of normalized signed distances. Next, distributions of MI values for male versus female neurons were derived as the marginals from the scatter plots for both male and female attention. The histogram for each group was fitted using a nonparametric kernel distribution to represent the probability density function (pdf) of the quantity. Given the heavy-tail and non-normal trend of the distributions, a non-parameteric Mann-Whitney U test (MATLAB *ranksum* command) was used to compare differences between male and female neurons under both attentional tasks. For comparison, ferret responses from ferrets S and A were also analyzed using MI and differences between male and female neurons were compared following the same procedure defined above. Male and female neurons were selected either using SSI or CSI methods.

Finally, the model output for an example sound mixture was analyzed for a male/female mixture as well as the same mixture with the two voices shifted in frequency. This analysis probed the dependence of the model on strict labels in the frequency dimension and its ability to ‘attend’ to a voice regardless of the exact F0, harmonics and formant peaks. To contrast, we modified the model into a channel selection model where pre-attention embeddings *Mp* were processed differently by zeroing out all neurons that did not match the target and only leaving neurons that matched it. In the case of attention to the female voice, only female neurons were maintained while all male neurons were canceled. This modified embedding was then mapped back into an output spectrogram to evaluate the integrity of the reconstruction in the case of original or shifted speech.

## Acknowledgments

The authors would like to acknowledge funding from the National Institutes of Health (R01DC005779, R01DC016119 **NICDC** & 1U01AG058532 **NIBI**, and a training grant **T32**).

## Author Contributions

**Joshi Neha** worked on the design of Experiment I, single-unit recordings, analysis of the data, and writing of the relevant results in MS.

**Yu Ng** participated in single-unit recordings, data analyses, and editing of the MS.

**Karran Thakkar** worked on the computational modelling, its applications to the physiological data, and model description in the MS.

**Daniel Duque** participated in the design of the experiment, animal behavioral training, initial physiological recordings and data analyses.

**Pingbo Yin**, helped with the behavioral training of the animals and guidance during the physiological recordings.

**Jonathan Fritz** participated in the supervision of the experimental work and data analyses. Mounya Elhilali provided conceptual guidance to the experiments, and helped with data analyses and the computational modelling, and provided a detailed review of the manuscript at various stages.

**Shihab Shamma** worked on the design of the experiments, data analyses, and the writing of the manuscript.

